# Sahul’s megafauna were vulnerable to plant-community changes due to their position in the trophic network

**DOI:** 10.1101/2021.01.19.427338

**Authors:** John Llewelyn, Giovanni Strona, Matthew C. McDowell, Christopher N. Johnson, Katharina J. Peters, Daniel B. Stouffer, Sara N. de Visser, Frédérik Saltré, Corey J. A. Bradshaw

**Affiliations:** Global Ecology, College of Science and Engineering and ARC Centre of Excellence for Australian Biodiversity and Heritage, Flinders University, GPO Box 2100, Adelaide, South Australia 5001, Australia; Research Centre for Ecological Change, University of Helsinki, Viikinkaari 1, Biocentre 3, 00790, Helsinki, Finland; Dynamics of Eco-Evolutionary Pattern and ARC Centre of Excellence for Australian Biodiversity and Heritage, University of Tasmania, Tasmania 7001, Australia; School of Natural Sciences and Australian Research Council Centre of Excellence for Australian Biodiversity and Heritage, University of Tasmania, Private Bag 55, Hobart, Tasmania 7001, Australia; Centre for Integrative Ecology, School of Biological Sciences, University of Canterbury, Christchurch, New Zealand; Community and Conservation Ecology, Centre for Ecological and Evolutionary Studies, University of Groningen, PO Box 11103, 9700 Groningen, The Netherlands

**Keywords:** ecological network, coextinction, biotic interactions, extinction event, Late Pleistocene, food web

## Abstract

Extinctions stemming from environmental change often trigger trophic cascades and coextinctions. However, it remains unclear whether trophic cascades were a large contributor to the megafauna extinctions that swept across several continents in the Late Pleistocene. The pathways to megafauna extinctions are particularly unclear for Sahul (landmass comprising Australia and New Guinea), where extinctions happened earlier than on other continents. We investigated the role of bottom-up trophic cascades in Late Pleistocene Sahul by constructing pre-extinction (~ 80 ka) trophic network models of the vertebrate community of Naracoorte, south-eastern Australia. These models allowed us to predict vertebrate species’ vulnerability to cascading extinctions based on their position in the network. We tested whether the observed extinctions could be explained by bottom-up cascades, or if they should be attributed to other external causes. Species that disappeared from the community were more vulnerable, overall, to bottom-up cascades than were species that survived. The position of extinct species in the network – having few or no predators – also suggests they might have been particularly vulnerable to a new predator. These results provide quantitative evidence that trophic cascades and naivety to predators could have contributed to the megafauna extinction event in Sahul.

## Introduction

Of all the extinctions that have ever occurred on Earth, many — potentially most — have been coextinctions [1]. In some cases, these coextinctions involved host-specific parasites that were doomed by the extinction of their host species, or flowering plants imperilled by the extinction of their pollinators [2]. Coextinctions have also been mediated through trophic interactions between herbivores and vegetation, and between predators and their prey [3,4]. Therefore, if we are to understand past extinction events and predict future extinctions, we need to be able to infer coextinction cascades accurately.

Changes in the primary producer component of a community can trigger bottom-up cascades and profoundly alter ecological communities [5]. However, it is unclear which species are most vulnerable to bottom-up cascades. On the one hand, it has been argued that top predators and species from high trophic levels are particularly sensitive to food-web perturbations and reductions in habitat area/primary productivity [6–9]. However, others have concluded that changes in the diversity of primary producers or nutrient content most strongly affect herbivores, and the cascading effects on higher trophic levels are dampened by trophic distance [10–13]. Similarly, extinction-risk assessments by the International Union for Conservation of Nature (IUCN) and recorded recent extinctions suggest that herbivorous terrestrial vertebrates are particularly vulnerable to extinction [14], a pattern that might partly be explained by the sensitivity of lower trophic levels to bottom-up cascades. The uncertainty regarding how vulnerability to bottom-up cascades varies with species traits (such as trophic level) has limited our ability to assess the importance of bottom-up cascades in past extinction events, and to predict how these cascades might unfold in the future.

Identifying the vulnerability of species to bottom-up (or top-down) coextinction cascades relies heavily on understanding species interactions within an ecological community. To this end, ecological network modelling is an invaluable tool for representing ecological communities from the perspectives of species interactions and for studying the consequences of changes in these interactions [15]. In ecological network models, organismal groups (e.g., species, age groups, populations, or individuals) are represented by nodes, and interactions — which can be weighted or unweighted — are represented by links (edges). The interaction type most frequently used to build ecological network models are trophic interactions (i.e., food webs). For contemporary communities, there is a growing number of studies that use detailed information on species interactions to build network models and to study trophic cascades [16–18]. Unfortunately, similar approaches are challenging to apply to palaeo-communities because of the lack of data on ancient trophic interactions. However, by combining contemporary and palaeo-data to infer trophic interactions, this limitation can be overcome and network models of palaeo-communities can be constructed [19,20]. For example, Pires et al. (2015) used this approach to model Late Pleistocene mammal communities in the Americas, concluding that (1) pre-existing American mammal networks were not especially unstable (i.e., they were similar to modern networks in Africa in terms of population densities re-establishing after simulated perturbations), and (2) the arrival of humans destabilized the networks because this new predator increased network connectance (i.e., the proportion of potential links that are realized). Investigations of paleo mass extinctions are fortified through the use of network modelling, as these methods provide insights into the causes and consequences of extinction events. Such events can also be used as a means of validating modelling methods because the outcomes (i.e., extinctions) are known. Despite these opportunities, the application of network modelling to investigate palaeo-extinction events remains under-utilised.

Megafauna (animals > 44 kg) extinctions swept across several continents during the Late Pleistocene (126,000 – 12,000 years ago), with the highest proportions of genera lost from Sahul (landmass including Australia and New Guinea) and the Americas [21,22]. While the causes of these extinctions are still debated, most evidence points toward the arrival of anatomically modern humans and/or climate change [23–25]. Irrespective of the root causes, large extinction events such as these always involve both primary and secondary (or co-) extinctions [3]. Indeed, it has been argued that the loss of prey species led to large predators going extinct in the Late Pleistocene [26,27]. Although the arrival of modern humans and/or climate change have been identified as the most likely ultimate causes of megafauna extinctions in the Late Pleistocene, vegetation change associated with human arrival and/or climate change has been identified as a pathway through which these ultimate causes could have triggered extinctions (i.e., via bottom-up trophic cascades) [28–31].

The Late Pleistocene megafauna assemblage of Sahul was distinct from that of other continents in that all the large mammals were marsupials or monotremes [32]. Giant reptiles and birds were also a prominent component of the continent’s megafauna [33]. While Sahul’s megafauna included many species over the standard body-mass threshold of 44 kg, the term ‘megafauna’ is often extended to include species with a body mass above that of their surviving relatives [32] — a definition we have adopted here. Identifying the pathways by which Sahul’s unique megafauna were lost is challenging because their extinctions happened much earlier in Sahul than elsewhere [34]. To characterize such ancient extinction events, a sufficient number of dated fossilized remains is necessary [35]. The most detailed and well-studied fossil record spanning the megafauna extinction event in Sahul comes from the Naracoorte region in south-eastern Australia (Figure 1). This fossil record offers an exceptional picture of the species living in the region over the past 500,000 years, including the ecological community at the time of the main megafauna extinction event that occurred approximately 44,000 years ago in Naracoorte [25]. Thus, the Naracoorte fossil record is the best platform available from which to model the ecological and environmental processes potentially involved in megafauna extinctions in Sahul.

**Figure 1.**
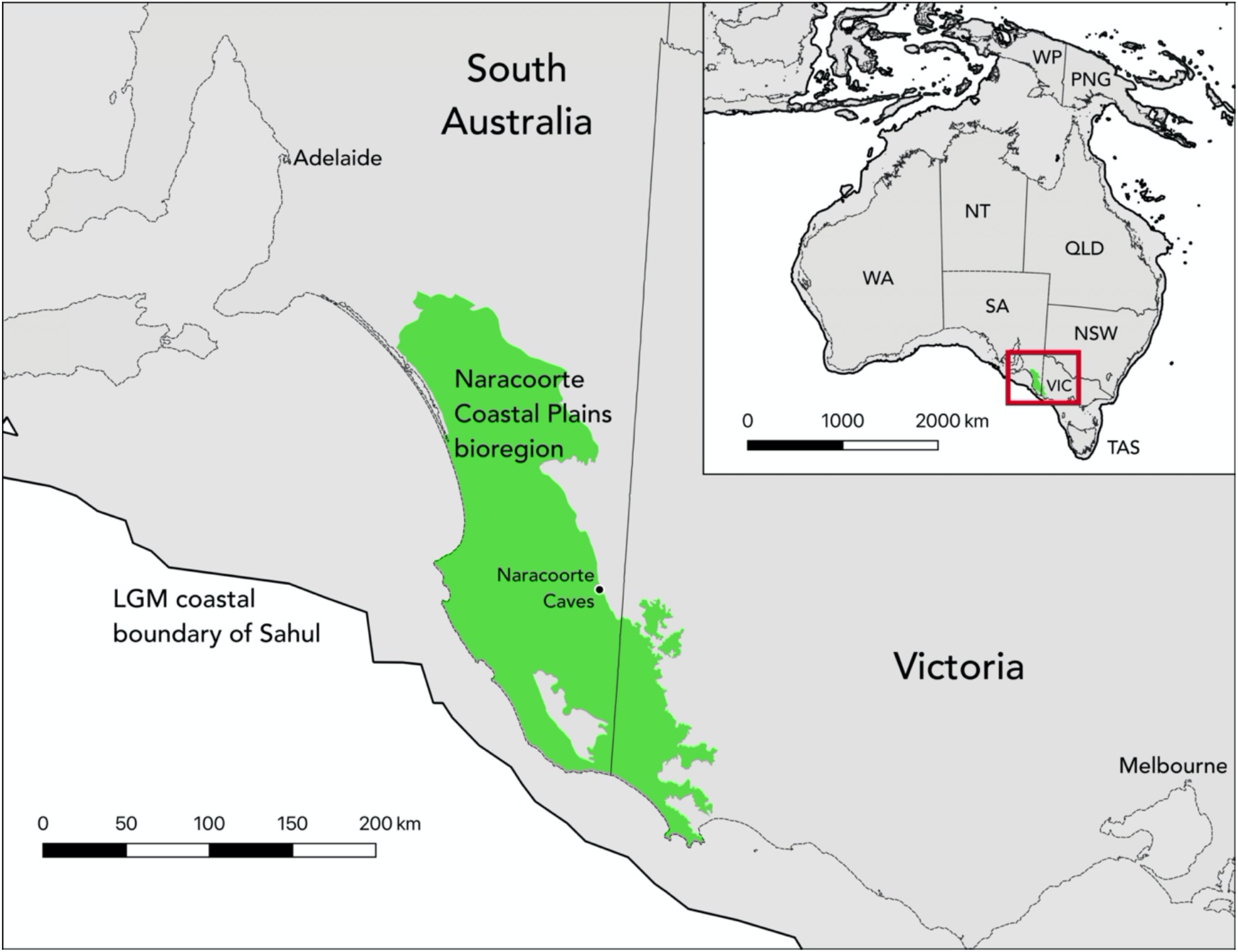
Sahul (top right insert) and the Naracoorte bioregion/Naracoorte Caves in south-eastern Sahul (main figure). These maps show coastline/sea level as they were during the Last Glacial Maximum (LGM; approximately 19,000 to 26,500 years ago). The grey area and thick outline indicate the land area during the LGM, the thinner lines show present-day coastlines and borders between countries and Australian states and territories, and the green area highlights the Naracoorte bioregion.

We assessed how vulnerability to bottom-up cascades varies with network-position attributes and whether bottom-up coextinction cascades stemming from the loss of basal resources (i.e., primary producers/plants) could have played a role in the megafauna extinctions of Sahul. First, we built small, synthetic networks (3 to 20 nodes) varying in topology (i.e., structure of connections), and calculated each node’s coextinction vulnerability using two methods: (*i*) simulation, and (*ii*) Bayesian networks. This allowed us to test the general influence of trophic level, diet breadth, and number of connected basal resources on vulnerability to bottom-up coextinction cascades.

Next, we used Naracoorte as a model system to assess whether bottom-up coextinction cascades could explain which species went extinct during the Late Pleistocene event. We constructed an entire terrestrial, palaeo-vertebrate assemblage (including all terrestrial vertebrate classes), and combined this assemblage with palaeo and contemporary data to infer trophic interactions and build network models (Figure 2). These network models consisted of nodes (species) with directed, unweighted links. We then computed each species’ vulnerability to coextinction via bottom-up cascades using the simulation method we validated with the synthetic networks, and we compared the coextinction vulnerabilities (and the traits influencing them) between extinct and extant (surviving into the Holocene) species. In addition to vulnerability to bottom-up coextinction cascades, we also tested for differences in the network positions of extant and extinct species to determine if relative position could have made extinct species more vulnerable in other ways (e.g., more vulnerable to a new predator). By incorporating interactions involving all vertebrate species, we adopted a holistic approach to studying megafauna extinctions of the Late Pleistocene.

**Figure 2.**
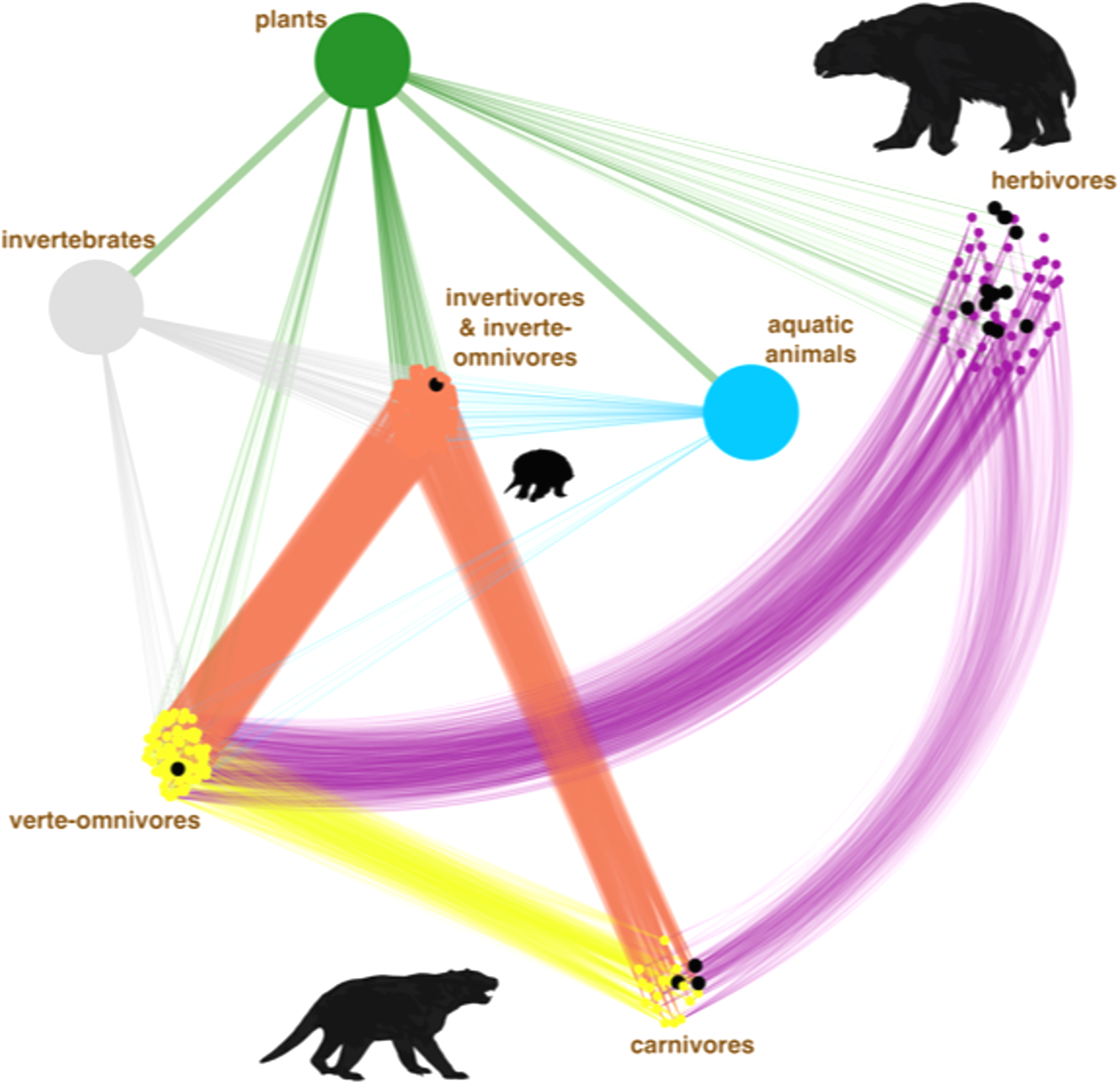
Example of an inferred ecological network model of the Late Pleistocene Naracoorte assemblage. Small points represent vertebrate species (nodes) and lines represent trophic interactions (links). Point colour shows trophic group (e.g., herbivores, carnivores etc), and extinct nodes are black. Due to uncertainty regarding trophic interactions, 1000 versions/models of the Naracoorte network were inferred and analysed. Plants, invertebrates and fish are shown as single large points in this figure.

We found that vulnerability to bottom-up cascades decreases with increasing trophic level, diet breadth, and basal connections, and that extinct species were more vulnerable to bottom-up coextinction cascades than were extant species. This suggests that bottom-up trophic cascades possibly contributed to the megafauna extinction event in Sahul. Our results also indicate that extinct species had fewer predators than did surviving species, suggesting that extinct species might have been particularly sensitive to the arrival of the new predator *Homo sapiens*.

## Methods

We aimed to: (1) identify how vulnerability to bottom-up cascades varies with trophic level, diet breadth, and basal connections using synthetic networks, and (2) develop and interrogate ecological network models representing the Naracoorte ecosystem prior to the main megafauna-extinction pulse and the arrival of humans in the region (~ 44,000 years ago) [25]. Below, we describe how we generated the synthetic networks, measured node vulnerability to bottom-up cascades, built ecologically realistic models of the Naracoorte network, and tested whether bottom-up coextinction cascades could explain which species were lost from the Naracoorte network (see Supplementary Figure S1 for a flowchart of methods for the construction and analysis of the Naracoorte network).

### Synthetic networks

We randomly generated 196 ‘synthetic’ networks that differed in topology. The networks varied in size (number of nodes: 3 to 20), number of links (2 to 92), and connectance (0.07 to 0.33). Within these networks, nodes varied in trophic level (1 to 4), number of ‘in’ links (1 to 8), and the number of basal nodes to which non-basal nodes were directly or indirectly connected through ‘in’ links (1 to 8). We generated these networks to test how a node’s vulnerability to the removal/extinction of basal nodes (analogous to plants/primary producers) varied depending on the node’s trophic level, diet breadth (number of ‘in’ links), and the number of basal resources to which it was connected.

### Coextinction vulnerability in synthetic networks

We inferred vulnerability of nodes to bottom-up cascades via: (*i*) simulations, and (*ii*) a Bayesian network method [36]. We applied two different approaches because there is uncertainty regarding the best methods to infer vulnerability to bottom-up cascades [3], so consensus using both approaches would provide more robust results than relying on only one. In the simulation method, primary extinctions occur by randomly removing basal nodes from the network, after which coextinctions are simulated by removing nodes that had lost all their ‘in’ links. Based on 1000 iterations of each network, we calculated the average coextinction vulnerability of each node as the proportion of total basal resources remaining when coextinction occurred. Applying the Bayesian-network method to the same networks [36], baseline extinction probabilities are assigned to each node and then each node’s accumulative extinction vulnerability is calculated using the network structure (taking into account each node’s dependencies on other nodes). In our case, we adjusted baseline probabilities of extinction so that primary extinctions were restricted to basal nodes.

### Analysis of synthetic networks

We fit mixed-effects models to the results from the simulation and Bayesian network approaches to assess if the two methods yielded similar results in terms of the effects of trophic level, diet breadth, and basal connections on vulnerability to bottom-up coextinction cascades. Prior to fitting, we scaled the independent variables so that the units of the regression coefficients were the same for all variables. The full/global mixed-effects model had coextinction vulnerability as the dependent variable (response), trophic level, diet breadth, number of connected basal nodes, and the interactions between these traits as independent variables (fixed effects), and network identity as a random effect. We compared support for the full models with every combination of nested/reduced model using Akaike’s information criterion weights (*w*AIC_*c*_). If including trophic level, diet breadth, and number of basal connections resulted in models with higher *w*AIC_*c*_, this suggests these variables affect vulnerability. We calculated model-averaged (via *w*AIC_*c*_) coefficients for the independent variables to identify how each variable influenced vulnerability. We also extracted marginal R^2^ from three mixed effects models that had either trophic level, diet breadth, or basal connections as the only independent variable to estimate how much variation in vulnerability each of these variables explained.

### Naracoorte study region

The World Heritage-listed (from 1994) Naracoorte Caves in south-eastern South Australia (37° 02′ 24″ S, 140° 48′ 00″ E) encompass a series of limestone caves that opened and closed to the surface at different locations, and at different times, over the last 500,000 years [37]. These openings acted as natural pitfall traps, capturing snapshots of Naracoorte’s biodiversity at different periods from at least 500,000 years ago to the present. In addition to the fossils of the many animals that fell into these natural traps, there are remains of species that lived in the caves, such as owls and bats, and their prey [38]. Consequently, the Naracoorte Caves provide an ideal platform from which to build palaeo-ecological network models to gain insight into how these long-lost ecosystems functioned and changed over time.

### Species data

To build a species assemblage list (i.e., to identify the nodes to include in the network models), we used two data sources: *FosSahul* 2.0 [35,39] and the Atlas of Living Australia. *FosSahul* 2.0 is a database of dated fossil records from Sahul, including an automated quality-rating method for date reliability [40]. We extracted and vetted records from the Naracoorte region (defined as the region between 35° 32′ 48″ S and 38° 6′ 50″ S, and between 139° 10′ 42′*’* E and 141° 0′ 21′*’* E) from *FosSahul* 2.0 that had high- or intermediate-quality dates (A*, A or B) [35], and whose age was younger than 200,000 years before present. We chose this cut-off age to provide a large enough period to include dated fossils from all/most megafauna species that lived in the region immediately prior to the main extinction event; if we had made the period too narrow, few megafauna species would have been captured despite their likely presence in the region at the time. Fossil records suffer from taphonomic biases (biases in the accumulation and preservation of different organisms), and, consequently, some species that were present in Late Pleistocene Naracoorte are unlikely to be represented in the fossil record. Furthermore, there are biases for studying and dating particular groups of species due to academic and amateur interests [41]. These biases, along with the fact that *FosSahul* was primarily designed to document megafauna remains rather than smaller species, means that *FosSahul* 2.0 does not include all vertebrate species present in Late Pleistocene Naracoorte. To account for this gap in the species list, we supplemented *FosSahul* 2.0 data with contemporary and historical species records from the Naracoorte Coastal Plains bioregion from the Atlas of Living Australia online repository (ala.org.au; accessed 3 January 2019). The Atlas of Living Australia has detailed species records of vertebrates from this bioregion, and so it captures most of the diversity of extant and recently extinct vertebrate species. We extracted data pertaining to all terrestrial vertebrates from the region, and removed species that were introduced since European arrival, as well as vagrants and erroneous records (species well-outside their normal distribution and not present in the fossil record), and strictly coastal species such as marine birds that do not use inland waterways. Our final species list, built using *FosSahul* 2.0 and the Atlas of Living Australia, included 280 birds, 81 mammals, 50 reptiles, and 12 amphibians (Supplementary Table S1).

The fossil record and phylogeography of extant mammal species (i.e., the vertebrate class most intensively studied) from the Naracoorte region suggest that, at a fine spatial scale, the distributions of some species expanded and contracted with climatic fluctuations during the Late Pleistocene and Holocene, but that during times of contraction species persisted at a regional scale (as in our study) in refugia [42–44]. Furthermore, most extant mammals recorded in the Late Pleistocene fossil record at Naracoorte were present (living) in the Naracoorte bioregion when Europeans arrived, suggesting little species turnover (apart from megafauna) between the Late Pleistocene and European arrival [42–44]. Together, these results suggest that our approach of using the fossil record and modern presence data provides a reasonable estimate of the species likely present in the region in the Late Pleistocene.

To infer trophic links, we required information on each species’ body mass and broad trophic category (whether it ate plants and/or fungi, invertebrates, terrestrial vertebrates, or fish). For extant species, we extracted much of this information from large databases: snake database [45]; Australian bird database [46]; *PanTHERIA* (mammal database) [47]; lizard database [48]; *AmphiBIO* (amphibian database) [49]; tropical bird database [50]; Amphibian database [51]; reptile database [52]; and Elton traits databases (mammals and birds) [53]. However, information on body mass was not available for some extant reptiles and amphibians. For these species, we estimated body mass using their body length (from field guides) and validated allometric relationships [45,48,51,54,55]. Extinct megafaunal species were not included in any of the aforementioned databases, so for these species we obtained body mass and diet data from the literature (see Supplementary Table S2). The 423 vertebrate species in the Naracoorte assemblage included 125 that consumed vertebrates, 249 that consumed plants, 362 that consumed invertebrates, and 48 that consumed fish (Supplementary Table S1). Of the 423 species, 273 consumed more than one of these resource groups. Mean species body mass ranged from 0.4 g to 2700 kg (Supplementary Table S1).

### Inferring trophic links

For almost all extant species, information on trophic interactions is incomplete, but the lack of knowledge regarding trophic interactions is even greater for long-extinct species. To overcome this limitation and build realistic ecological network models, various methods have been developed for inferring trophic links using species’ traits such as body size (i.e., larger predators tend to eat larger prey, and predators are usually bigger than their prey) [56]. Others have built on this approach to improve prediction accuracy [57–60] by adding phylogenetic and physiological information, prohibiting impossible or unlikely links, incorporating specific morphological features such as biting force and cuticular thickness, and taking into account abundance. However, most of this research focusses on fish or invertebrates, with the effectiveness of these methods rarely applied or validated for terrestrial vertebrates, but see [61,62].

We therefore developed and validated a new method, based on the body-size trophic-niche model [56], to infer trophic links between terrestrial vertebrates. The body-size trophic-niche model that we adapted consists of two quantile regressions: (*i*) one defining the upper prey-size limit given predator mass, and (*ii*) the other defining the lower prey-size limit given predator mass. If a species falls within the upper and lower limits for a particular predator, it is inferred as potential prey for that predator. We used a large predator-prey interaction dataset to identify these body-size relationships between terrestrial predators and their prey, and tested whether these relationships varied depending on the predator’s taxonomic class (i.e., did including predator class as an independent variable improve the fit of the body size quantile regressions?).

We extracted the interaction dataset from *GloBI*, an online repository of biotic interactions [63]. The dataset consisted of 3893 records: 958 records of predation by non-marine mammals, 2711 by birds, 199 by reptiles, and 25 by amphibians (Supplementary Table S3) [64]. We extracted data on mean body mass for the species from the same databases we used to add this information to the Naracoorte species list (see above). Once we identified the best trophic-niche model (i.e., the combination of quantile regressions that best fit the upper and lower prey-size limits) using the entire *GloBI* dataset and Bayesian information criterion (BIC; Supplementary Methods S1), we validated this method of assigning trophic links by comparing model performance when applied first to the *GloBI* data divided into training and validation datasets, and then to a well-resolved trophic network from the Serengeti [65] (see also S. de Visser unpublished data; Supplementary Methods S1). We used the true skill statistic to evaluate model performance and found that the top-ranked model (according to BIC) also performed best at assigning links in the validation step (Supplementary Methods S1, Supplementary Table S4, Supplementary Figure S2, script available at [64]). We used the best trophic-niche model to identify potential prey for each predator of vertebrates (see methods below; Supplementary Methods S1; script available [64]).

While including additional species traits could improve the accuracy of inferred predator-prey interactions, we used only three readily available traits (broad diet, body size, and predator taxonomic class — with reptiles and amphibians grouped together). We used only these three traits because: (1) using few traits is compatible with the quantile regression framework, (2) they can easily be extracted for vertebrate species in most assemblages (and therefore the method can be widely applied), and (3) our validation steps demonstrated the resulting performance of the trophic-niche model is sufficient (true skill statistic [TSS] = 0.6 when applied to the Serengeti assemblage; TSS varies from −1 to 1, with a score of 0 indicating no better than random; Supplementary Table S5).

### Naracoorte networks

To build realistic ecological networks for the Naracoorte assemblage, we applied the trophic-niche model to the species list, removed excess links between vertebrates (to account for overestimating the number of predator-prey links), and added links to non-terrestrial vertebrate food resources. However, there is uncertainty regarding which vertebrate predator/prey links to delete as well as how many links to add from non-terrestrial vertebrate food resources to vertebrates. To address this uncertainty, we used a randomization approach in the link-removal and -addition steps described below, and generated 1000 versions of the network. That is, we randomly removed (for the vertebrate predator-prey links) or added (for the herbivores, invertivores, and piscivores) links in the range indicated as realistic based on contemporary species’ diet breadths. Because we do not know exactly where to add or remove these links, we generated 1000 versions of the network so our results were not skewed by the particular links selected.

We used the trophic-niche model to assign potential predator-prey links in the Naracoorte species assemblage. Although trophic-niche models are good at identifying potential links, they almost always overestimate the number of realized links [66]. This is because predators are unlikely to consume all prey within their size range — some species are not palatable, are dangerous, too rare, difficult to capture, use different microhabitats, or have other ecological characteristics that make them unsuitable for regular consumption [66]. To build a network with a more realistic structure, we assigned a probability to each interaction based on the prey’s position in the predator’s prey-size range and a Gaussian distribution centred on this range (with a standard deviation equal to one quarter of the predator’s prey-size range); the highest probability was for prey close to the centre of the prey-size range (i.e., centre of the distribution) and decreased the closer the prey was to the predator’s limits. For each predator, we randomly sampled from a density kernel fit to published carnivore diet breadths (*n* = 12; sampled between 1 and twice the maximum diet breadth in Supplementary Table S6a; Supplementary Methods S1) to select the number of potential prey that were ‘realized’ prey. In assigning the sampled diet breadths to individual predators, predators with more potential prey (indicated by the trophic-niche model) were assigned larger diet breadths than were those with fewer potential prey. To account for different degrees of dietary specialisation, we Poisson-resampled the number of potential prey for each predator before ranking predators according to their number of potential prey, slightly shuffling relative diet breadths between network models. Then, using the assigned diet breadths, we randomly selected from the potential prey, taking into account the probability of the predator-prey interaction. This method resulted in a vertebrate network with realistic connectance (proportion of potential links that are realized), and with most of each predator’s prey closer to the centre, rather than the limits, of their prey-size range.

Terrestrial vertebrates not only consume other terrestrial vertebrates, they also consume invertebrates, plants, fungi, and fish. In addition to inferring trophic links among terrestrial vertebrates, we therefore needed to add links to vertebrates from these other food resources. However, we did not include detail on individual species within these resource groups because: (*i*) our study focusses on terrestrial vertebrate species; (*ii*) invertebrate, plant, and fungal diversities are not well resolved for most ecosystems (including for Late Pleistocene Naracoorte); and (*iii*) fish only constitute a small part of the Naracoorte community in terms of biodiversity and biomass. We therefore generated a pool of *n* species for each of these groups (*n* = 1300 for plants, 6000 for invertebrates, and 23 for freshwater fish), with the number of invertebrate and plant species calculated based on the described diversity in these groups relative to terrestrial vertebrate diversity in Australia [67], and the number of fish determined by the diversity of freshwater fish recorded in the Naracoorte Coastal Plains bioregion in Atlas of Living Australia (ala.org.au; accessed 6 April 2021). To assign links to vertebrates from species in these groups, we used published records of vertebrate diet breadth for 20 herbivores, 6 invertivores, and 9 piscivores (Supplementary Tables S6a and 6b). We fit kernel densities to the invertivore, herbivore, and piscivore diet breadth data, and randomly sampled (within a diet breadth range of 1 to twice the maximum diet breadth recorded for that trophic guild in Supplementary Table S6a and S6b) from these distributions to assign the number of links between each resource group and vertebrate consumer in each of the 1000 network models. However, because the published piscivore diet breadths (Supplementary Table S6b) came from terrestrial Australian predators that are not exclusively piscivorous, we multiplied the number of fish in the diets of these predators by the number of resource groups from which the predator fed before fitting and sampling from the kernel densities. This modification was made to offset the adjustment accounting for inflated diet breadths in omnivores (described below).

For vertebrates that fed from more than one group (i.e., omnivores, which represent over half the vertebrates in this network), we proportionally adjusted the number of ‘in’ links (food resources) depending on from how many food groups they fed. For example, if a species consumed from two groups (e.g., vertebrates and invertebrates), we randomly deleted half of the links from each group; if they fed from three groups, we randomly removed two-thirds of the links from each food group (and so on). We made these deletions to prevent omnivores from having inflated diet breadths.

In some cases involving water birds, we used empirical evidence to avoid assigning unrealistic trophic links. For example, pelicans are large and, consequently, the trophic-niche model predicts that pelicans take large vertebrate prey. However, we know that pelicans are not birds of prey and do not consume large terrestrial animals. Thus, we restricted the allocation of trophic links for such birds to prevent them from feeding on unrealistically large terrestrial vertebrates (they were prevented from consuming prey that weighed over a third of their body mass).

We completed the networks by generating links from plants to invertebrates and from plants to fish. To determine diet breadth for these herbivores, we sampled from a Pareto distribution (alpha = 1.02, truncated at 52) following ref [68]. The alpha and truncation values were based on a temperate woodland system (i.e., similar to Naracoorte) [68–70].

### Analysis of Naracoorte models

We estimated the coextinction vulnerability of Naracoorte vertebrates using the same simulation approach as we applied to the synthetic networks i.e., plant nodes were iteratively (and randomly) removed to simulate primary extinctions, and coextinctions were triggered when animals lost all their food resources. We measured each species’ coextinction vulnerability as the proportion of plant nodes remaining when coextinction occurred. We repeated the simulations 1000 times for each of the 1000 network models to test whether: (1) the effects of trophic level, diet breadth, and basal connections (direct and indirect) on coextinction vulnerability were consistent with results obtained from the synthetic models, and (2) vulnerability to bottom-up cascades differed between extinct and extant species.

To test whether the coextinction vulnerability results were consistent with those from the synthetic models, we fit linear-regression models to the data with vulnerability to coextinction cascades as the response, and trophic level, diet breadth, basal connections, and the interactions between these variables as independent variables (fixed effects). Rather than using the raw data from the 1000 network models, we used each species mean value for each trait and vulnerability for these analyses. We then compared relative model probabilities (*w*AIC_*c*_) for the full model to all possible reduced models, and examined the coefficients of the main effects to determine if they were similar to those obtained from the synthetic networks.

We compared susceptibility of Naracoorte’s extinct *versus* extant species to bottom-up coextinction cascades in three steps. First, we compared the groups in terms of their trophic level, diet breadth, and number of basal connections to determine if differences in these variables suggest one group would be more vulnerable than the other. Second, to test for an association between coextinction vulnerability and extinction status, we compared *w*AIC_*c*_ support for two models: a null model with vulnerability to coextinction as the dependent variable and no fixed effects *versus* a model that was identical to the first model, except with extinction status as an independent variable. Third, we ran a randomization test to assess the probability that extinct species were more vulnerable to coextinction than were extant species. Here, we sampled the coextinction vulnerabilities of 10 extinct and 10 extant species from each of the 1000 models 20 times (i.e., using the raw data rather than species means), and each time calculated the mean difference in coextinction vulnerability between the two groups.

We also assessed the position of extinct *versus* extant species in the network using 12 different network metrics: *trophic level, pageRank*, *betweenness centrality*, *eigenvector centrality*, *closeness centrality (in)*, *coreness (in)*, *degree (in)*, *eccentricity (in)*, *closeness centrality (out)*, *coreness (out)*, *degree (out)*, and *eccentricity (out)* (Supplementary Table S7 for metric descriptions). We calculated the metrics followed by an ‘in’ or ‘out’ for each node using links pointing towards (in) or away from (out) the focal node. We chose these metrics because they are commonly used, node-level metrics describing position in the network. For each metric, we calculated the species’ mean value across the 1000 network models. After checking for highly correlated metrics and removing those identified as redundant (i.e., metrics that had a Spearman’s *ρ* > 0.8), we ran a principal component analysis and visually inspected for grouping of extinct and extant species. The reduced list of metrics included *closeness centrality (out)*, *eccentricity (out)*, *degree (in)*, *coreness (in)*, *betweenness*, and *PageRank*.

## Results

### Synthetic networks

The two methods calculating bottom-up coextinction vulnerability (simulation and Bayesian network) yielded similar results in terms of the effects of trophic level, diet breadth, and basal connections on node vulnerability to bottom-up coextinction cascades. *w*AIC_*c*_ indicated that the full models (i.e., that had all three independent variables and their interactions) were more strongly supported than reduced models using both approaches, with the full models having *w*AIC_*c*_ > 0.999 (Supplementary Tables S8 and S9). The weighted model-averaged coefficients describing the relationships between the three independent variables and coextinction vulnerability were negative, irrespective of which of the two methods we used to calculate vulnerability (Figure 3A; Supplementary Tables S8 and S9). These negative correlations indicate that vulnerability to bottom-up cascades decreased with increasing number of basal connections, diet breadth, and trophic level. Marginal R^2^ of the three reduced models, each of which had one of the three variables as a main effect, suggest that the number of basal connections explained most of the variation in coextinction vulnerability in the synthetic networks (Figure 3B). We restricted our analyses of the Naracoorte network models to the simulation method because both approaches yielded similar results, and because the Bayesian network method was prohibitively time consuming for networks of the size of the Naracoorte models.

**Figure 3.**
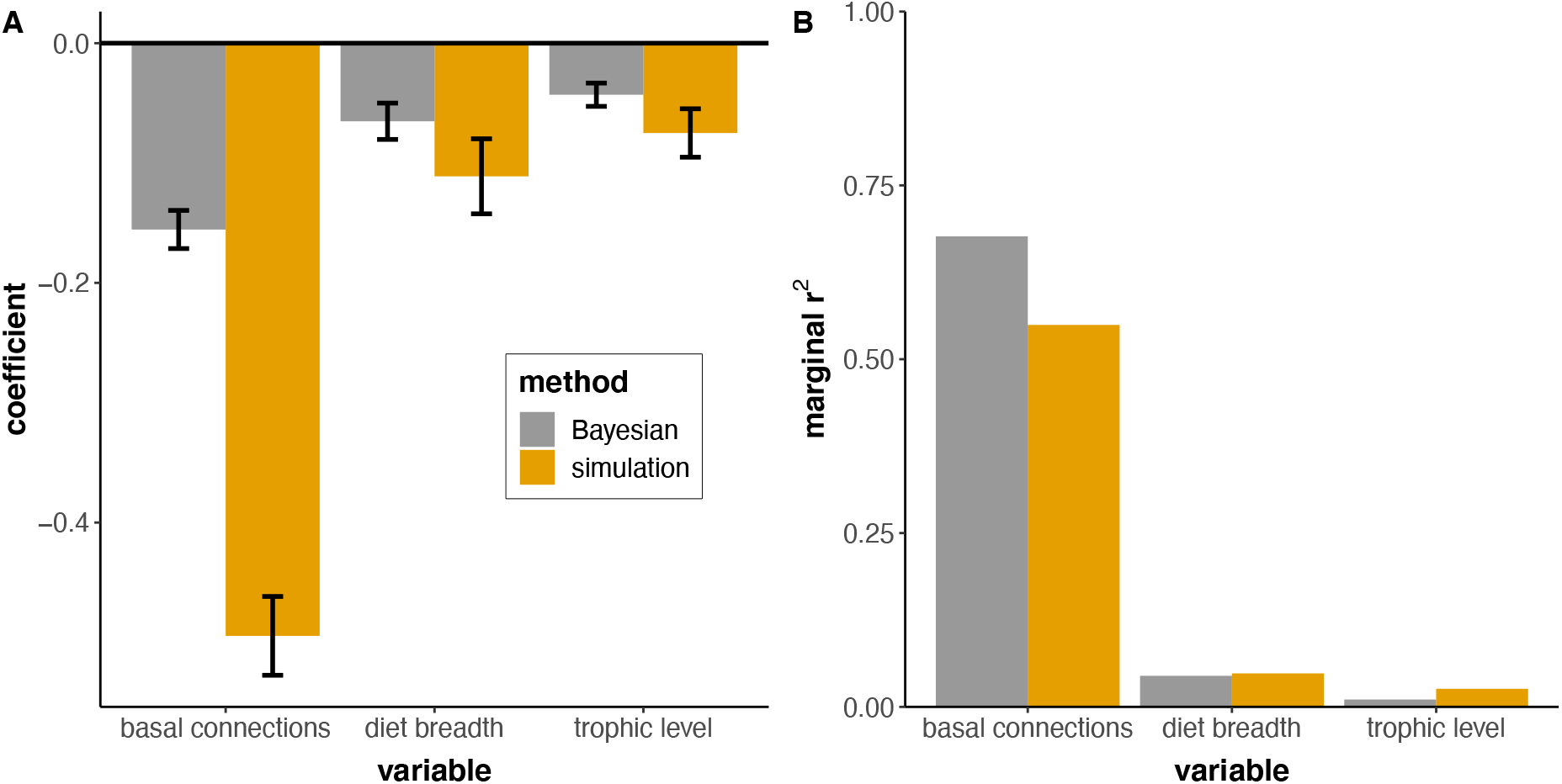
The effects of basal connections, diet breadth, and trophic level on vulnerability to bottom-up cascades in synthetic networks. Vulnerabilities were calculated using either a simulation or Bayesian network approach. Plot **A** shows the weighted, model-averaged coefficients of the main effects (both methods), and plot **B** shows the marginal R^2^ from mixed-effects models that had either basal connections, diet breadth, or trophic level as the only fixed effect (both methods). Error bars in plot A indicate 95% confidence intervals.

### Naracoorte network

In terms of the effects of trophic level, diet breadth, and basal connections on vulnerability, the patterns in the Naracoorte network were similar to those from the synthetic networks (Supplementary Table S10). *w*AIC_*c*_ strongly supported the full model over reduced models (*w*AIC_*c*_ ≈ 1 for the full model; Supplementary Table S10), and the three main effects were negatively correlated with coextinction vulnerability (Figure 4A; Supplementary Table S10). Reduced models that had either trophic level, diet breadth, or basal connections as the only independent variable had R^2^ > 0.23, indicating that each of these variables were associated with a substantial proportion of variation in vulnerability (Figure 4B).

**Figure 4.**
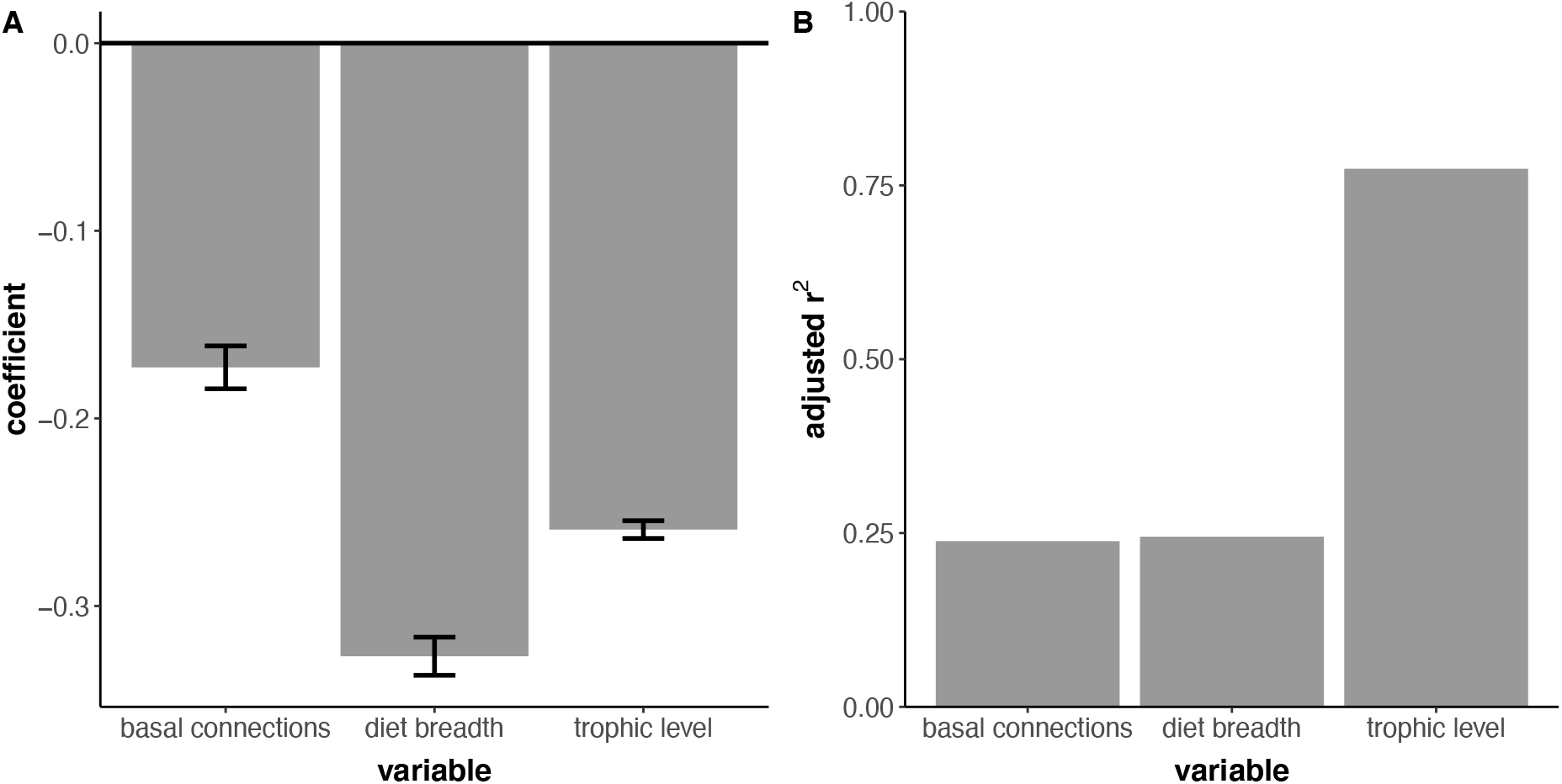
The effects of basal connections, diet breadth, and trophic level on vulnerability (calculated using the simulation approach) to bottom-up cascades in the Naracoorte network models. Each species’ mean number of basal connections, diet breadth, trophic level, and vulnerability to bottom-up cascades was calculated, and linear regression models were fit to the mean data. Plot **A** shows the weighted, model-averaged coefficients of the main effects, and plot **B** shows the adjusted R^2^ from linear regression models that had either basal connections, diet breadth, or trophic level as the only fixed effect. Error bars in plot A indicate 95% confidence intervals.

Extinct species had fewer basal connections, narrower diet breadths, and came from lower trophic levels, on average, than those species that survived into the Holocene (Figure 5A–C; mean basal connections: 298 *vs.* 515; diet breadth: 34 *vs.* 79; and trophic level: 2.3 *vs.* 2.9 for extinct *versus* extant species). Extinct species also had higher coextinction vulnerability than did surviving species (Figure 5D; mean coextinction vulnerability ± 95% confidence interval: 0.045 ± 0.015 *vs.* 0.008 ± 0.002 for extinct and extant species, respectively), a result consistent with coextinction vulnerability being higher for species with fewer basal connections, narrower diet breadth, and from lower trophic levels.

**Figure 5.**
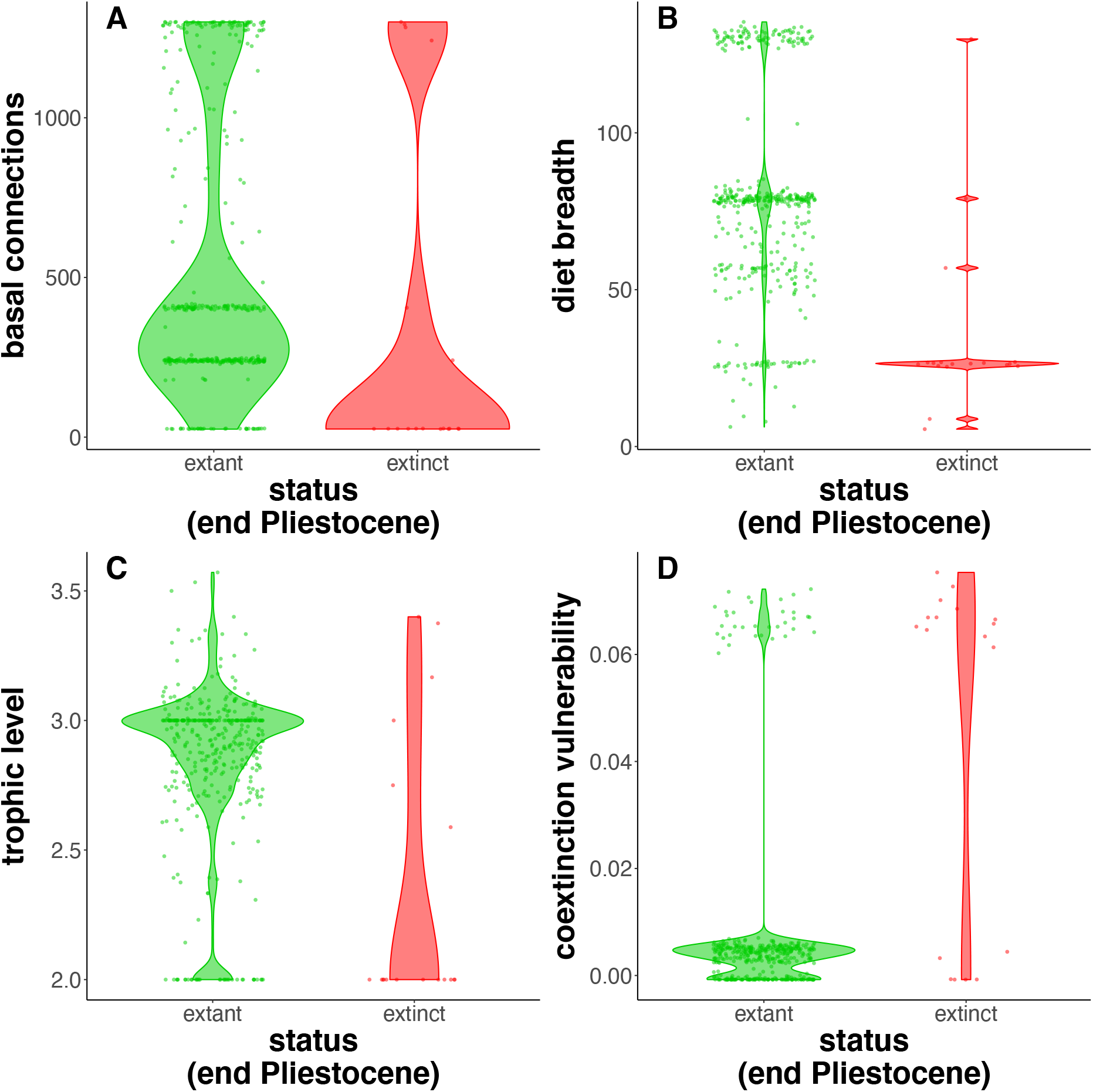
Comparison of species that survived into the Holocene (extant, shown in green) to those that went extinct in the Late Pleistocene (extinct, shown in red) from the Naracoorte network. Panel **A** shows the number of basal nodes (plants) connected directly and indirectly to each node via ‘in’ links; panel **B** shows diet breadth of each node; panel **C** indicates trophic level; and panel **D** shows the calculated coextinction vulnerability. The panels, which all include density violin plots and scatterplots, are based on each species’ mean score across the 1000 Naracoorte network models.

To test for an association between species’ extinction status and vulnerability to bottom-up cascades, we compared support for a model with coextinction vulnerability as the dependent variable and extinction status as the only fixed effect *versus* a null model (a random-intercept model with no fixed effects), and we also did a randomization test comparing the coextinction vulnerability of extinct *versus* extant species. Both approaches indicated that extinction status was associated with vulnerability to bottom-up cascades, with extinct species more vulnerable than those that survived into the Holocene. The *w*AIC_*c*_ for the model with extinction status as a fixed effect was ~ 1, indicating that this model was strongly supported over the null model. The randomization test showed that extinct species had a probability of 0.91 of being more vulnerable to bottom-up coextinctions than extant species (Figure 6).

**Figure 6.**
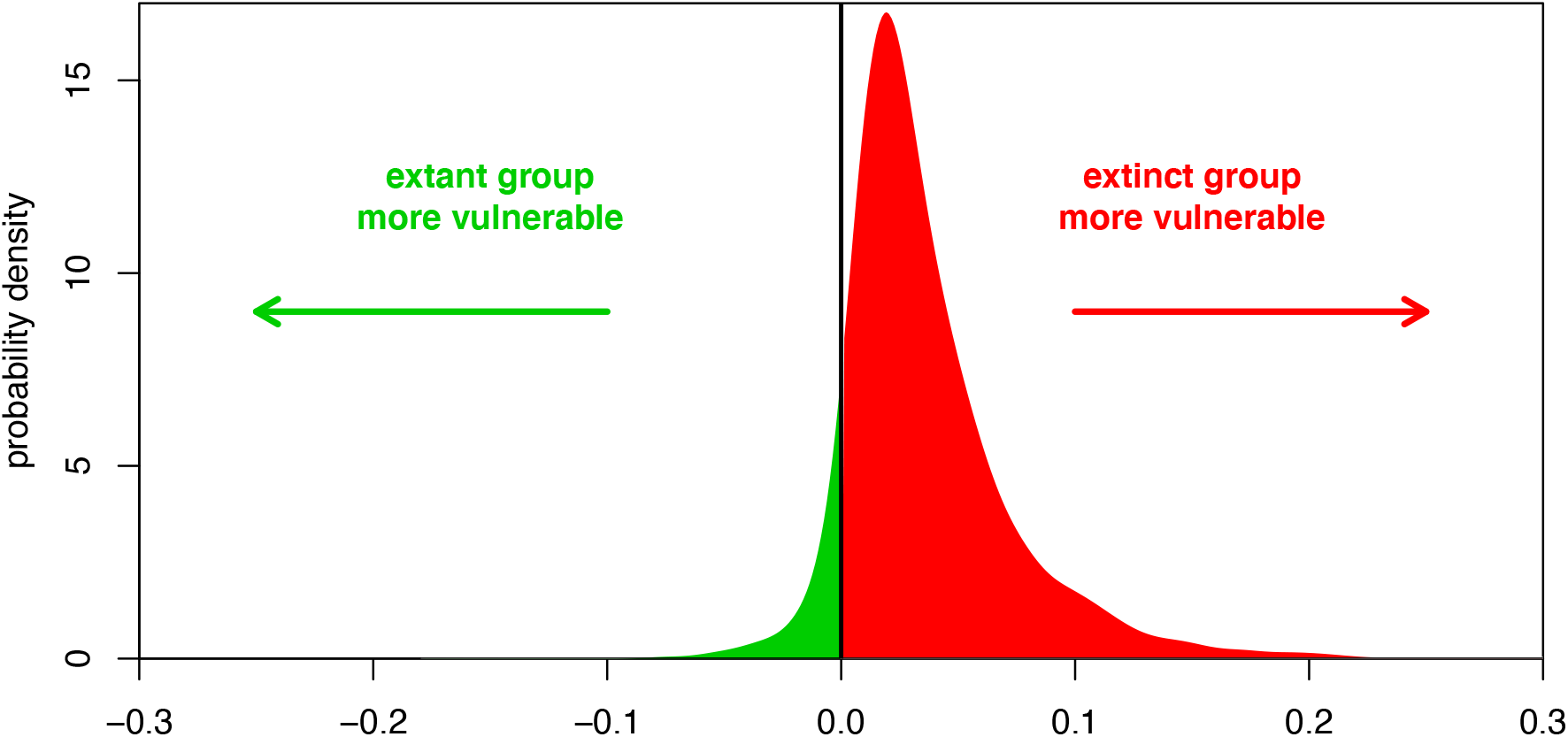
Density plot showing results from a randomization test (20,000 replicates) comparing bottom-up coextinction vulnerability of extinct *versus* extant species from the Naracoorte network models. Each species’ coextinction vulnerability was calculated using the simulation method (removing plant nodes and then removing animal nodes that no longer had connections to plant nodes). From these results, the coextinction vulnerabilities of 10 extinct and 10 extant species were sampled (with replacement), and the mean differences in coextinction between the groups calculated. This process was repeated 20 times for the results from each of the 1000 network models and used to build the density plot. The red area of the density plot indicates higher vulnerability for extinct species, and the green area indicates higher vulnerability for extant species.

Extinct species differed from extant species in terms of their position in the network. Principal component analysis of six network-position metrics showed that extinct and extant species fell into two distinct groups according to the second principal component (dimension 2 in Figure 7A). The main contributors to this principal component are metrics describing a node’s connection to the network through its ‘out’ links, including the closeness centrality (out) and eccentricity (out) metrics (Figure 7A; Supplementary Figures S3 and S4). Closer examination of the out links (i.e., number of predators) showed that, on average, extinct species had < 1 predator, whereas extant species had > 3 (Figure 7B; mean number of predators: 0.2 *vs.* 3.3 for extinct *vs.* extant species). Indeed, the average number of predators was lower for extinct than extant species in all 1000 models of the Naracoorte network.

**Figure 7.**
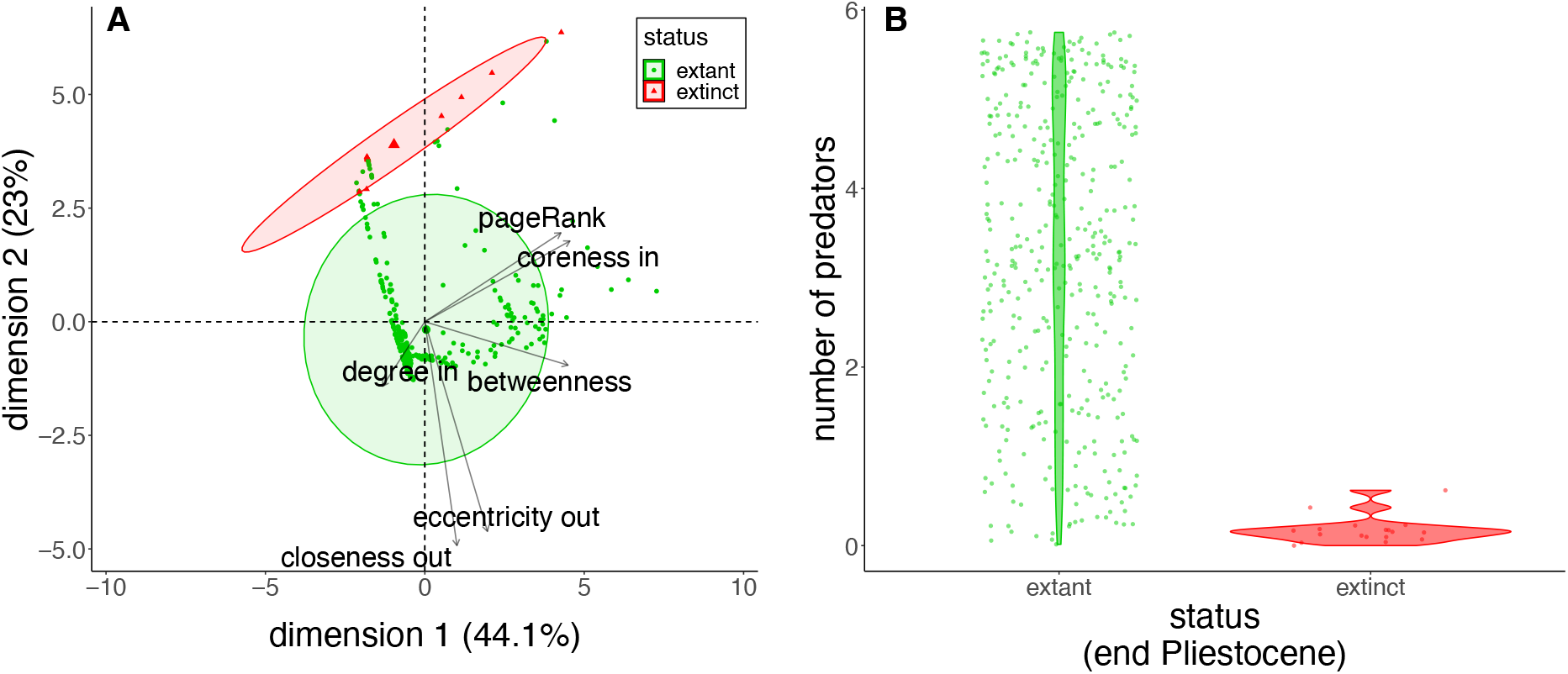
Comparison of extinct *versus* extant species in terms of their position in the Naracoorte trophic network. Panel **A** is a biplot of the first two principal components from a principal component analysis of six variables describing species’ positions in the network. Panel **B** is a violin and scatter plot showing the number of species that preyed on each extant *versus* extinct species. We inferred 1000 models of the Naracoorte network, and from these models calculated each species’ mean value of each metric for use in the principal-components biplot and predator-diversity plot.

## Discussion

Our analyses demonstrate that a species’ vulnerability to bottom-up coextinction cascades varies depending on its trophic level, diet breadth, and number of basal connections. We also found that the position of extinct species in the Naracoorte network — being primarily herbivorous (low trophic level), and therefore having relatively narrow diet breadths and few pathways to basal resources — might have made them more vulnerable to bottom-up coextinction cascades than were co-occurring species that survived into the Holocene. The Naracoorte results suggest that trophic cascades potentially contributed to the megafauna extinction event in south-eastern Sahul. However, the extinct species from Naracoorte also had fewer predators than did extant species, a network position attribute that would likely have made them more vulnerable to the arrival of the new ‘super predator’ — humans [71].

The Naracoorte and synthetic network models revealed that vulnerability to bottom-up coextinction cascades precipitated by plant extinctions decreased with increasing trophic level, diet breadth, and number of basal connections (Figures 3 and 4; Supplementary Tables S8, S9 and S10). Our results therefore support previous findings that species with narrower diet breadths/fewer pathways to basal resources are more vulnerable to bottom-up coextinction cascades [72–74]. However, previous research does not provide a clear expectation regarding the relationship between trophic level and extinction vulnerability. It is often assumed that higher trophic levels are more vulnerable to extinction than are lower levels due to the cumulative effects of disturbances on lower trophic levels (on which higher trophic levels depend), and direct persecution by humans [8,9]. However, our analyses that specifically tested for sensitivity of species to primary extinctions in the plant community imply that vulnerability to these bottom-up cascades in fact *decreases* with trophic level.

Consistent with these results, several manipulative experiments of ecological communities have revealed that changes in the plant component of the community most strongly affect herbivores, and impacts on higher trophic levels diminish with trophic distance [10,13,75].

This pattern has also been identified in theoretical studies. Applying Rosenzweig– MacArthur models and synthetic (but ecologically feasible) networks, the loss of primary producers triggered extinctions in herbivores before doing so in carnivores, and herbivores were more vulnerable to these cascades than were carnivores [74]. However, our vulnerability scores were based on coextinction being triggered when a consumer lost all food resources. Coextinctions could be triggered at lower thresholds and/or vary between species. While the congruence between our results and those from previous studies support the methods and threshold we used, further investigation into how coextinction threshold influences the effect of network position on node vulnerability, as well as how coextinction thresholds covary with species/community traits, is needed to refine methods for predicting the probability and magnitude of bottom-up cascades.

The extinction pattern observed in the Naracoorte assemblage could have been the result of bottom-up cascades triggered by changes in the plant community, as demonstrated by our vulnerability modelling. This leads to the question: did vegetation change at the same time as the megafauna disappeared? Studies in other regions of Sahul have detected shifts in vegetation roughly coinciding with, or immediately preceding, megafauna extinction. Hypothesised drivers of these vegetation shifts include land-use changes associated with human arrival (i.e., fire-stick farming) [28,76] and/or climate change [77]. However, there are no detailed studies on the vegetation of Naracoorte spanning the Late Pleistocene extinction event (but see [78] for a review of broad proxies of vegetative change over this period). The megafauna’s disappearance from Naracoorte did, nonetheless, coincide with an unusually cool period (Supplementary Figure S5) and the arrival of humans (~ 44,000 years ago) [25], offering the intriguing possibility that changes in climate and/or land use triggered shifts in vegetation that had consequences for higher trophic levels in the network. To evaluate this possibility, more studies are required to model vegetation changes in south-eastern Sahul (including the Naracoorte region), and these must be validated using the pollen record and/or other fossil data.

By considering the network position of all vertebrate species in the assemblage, a clear difference between extinct and extant species emerged — extinct species had fewer predators than did species that survived (mean number of predators: 0.2 *versus* 3.3 for extinct *versus* extant species, respectively; Figure 4B). This predator naivety, coupled with the species’ slow life histories, likely made megafauna especially vulnerable to new predators [34,79–81] and suggests that hunting by humans could have adversely affected megafauna. Thus, a network modelling approach to assessing extinction vulnerability suggests that bottom-up and/or top-down processes could have selectively removed the now-extinct species from the Naracoorte community. However, there remains scope to address uncertainties regarding the structure of the Naracoorte network and the methods for estimating vulnerability to ecological cascades. As palaeo-vegetation, invertebrate, trophic (including detailed information on the diets of extinct species), and climate data improve, network models can incorporate this information to build more refined networks, and include more detailed top-down and bottom-up forcings to assess the plausibility of the different potential causes and pathways to extinction — including what (if any) role humans played in the megafauna’s demise.

In summary, our network modelling of Late Pleistocene Naracoorte suggests bottom-up coextinction cascades and/or predator naivety and the arrival of humans could have contributed to the megafauna extinction event in Sahul. Indeed, that our network models showed that extinct species were vulnerable to bottom-up cascades *and* new predation pressures lends support to recent research suggesting that climate change (that can shift vegetation and lead to bottom-up cascades) and human arrival together drove the megafauna extinction trajectories in much of south-eastern Sahul [25]. Our research, along with other recent studies [19,62,82], demonstrates that network modelling is a powerful tool for investigating and understanding ancient extinction events. By developing methods to model whole-community responses to change and validating these methods using ancient extinction events, we can also provide better estimates of the fates of contemporary communities as the sixth mass-extinction event unfolds [83].

## Acknowledgements

We thank E. Reed, C. Carbone, M. Tucker, C. Dickman and V.K. Llewelyn for their constructive input.

