## Supplementary text, figures, and tables for "Sahul’s megafauna were vulnerable to plant-community changes due to their position in the trophic network"

**Supplementary Methods S1**

**Validation methods for trophic niche models**

To assign links between nodes (species), we used trophic niche-space models (e.g., [1]). Each of these models has two quantile regressions that define the prey-size range a predator of a given size is predicted to consume. Species whose body mass is within the range of a predator's prey size, as identified by the trophic niche-space model, are predicted to be prey, while those outside the range are predicted not to be eaten.

The broad taxonomy of a predator helps to predict predation interactions [2]. To optimize our trophic niche-space model, we tested whether including taxonomic class of predators improved the fit of quantile regressions. Using trophic (to identify which species were predators), body mass, and taxonomic data, we fitted and compared five quantile regression models (including a null model) to the *GloBI* data. In each model, we $\log_{10}$-transformed the dependent variable *prey body mass*, and included for the independent variables different combinations of $\log_{10}$-transformed *predator body mass*, *predator class*, and the interaction between these variables (Supplementary Table S4). We $\log_{10}$-transformed both predator and prey body mass to linearize the relationship between these variables. We fit the five quantile regressions to the upper and lower 5% of prey body mass, and compared model fits using the Bayesian information criterion (BIC). The *predator body mass*predator class* model fit the $95^{th}$ quantile data best, whereas the *predator body mass + predator class* model fit the $5^{th}$ quantile data marginally better than the aforementioned interaction model (Supplementary Figure S2, Supplementary Table S4).

Next, we compared the performance of two trophic niche-space models using empirical data and the true skill statistic (i.e., model validation). The true skill statistic assesses how well a model predicts present and absent links [1,3]:

$$\frac{(ad - bc)}{(a + c)(b + d)}$$

where ***a*** = number of predicted links observed, ***b*** = number of predicted links not observed, ***c*** = number of links observed but not predicted, and ***d*** = number of links predicted to be absent and not observed.

The two trophic niche models we compared were: (1) the simple body-size model originally developed by Gravel et al. [1], and (2) our new model based on the quantile regressions with the lowest BIC (i.e., *predator body mass*predator class* for the upper limit and *predator body mass + predator class* for the lower limit). We applied these trophic niche models to two empirical datasets for validation. In the first validation, we split the *GloBI* interaction data into training and validation datasets; we randomly assigned 75% of the interactions in the *GloBI* interaction dataset to the training dataset, and 25% to the validation dataset. Because *GloBI* only includes observed predator-prey interactions, we needed to generate 'observed absent' links to calculate the true skill statistic. We generated these absent links by randomly selecting pairs of species in the validation data that were not observed in trophic interactions with each other, making the same number of these absent links as there were observed links (1:1 ratio). We randomly generated 100 training and validation datasets in this way, and recorded the performance (true skill statistics) of the two trophic niche models on these sets.

The second dataset we used to validate our trophic niche models was a real-world food web from the Serengeti (de Visser et al., 2011, S. de Visser, unpublished data). This is probably the most highly resolved, diverse terrestrial vertebrate food web that has been documented and published; it describes one of the most intact ecosystems today, and one that includes much of its Late Pleistocene megafauna (i.e., it is more analogous to Late Pleistocene assemblages in Australia in this regard). While it is the most detailed diverse terrestrial vertebrate food web in existence, it consists of functional/trophic species groups rather than individual species. We ran the validation analyses on the Serengeti food web in the same way as described for the *GloBI* training/validation datasets, only in this case we: (1) used the full *GloBI* dataset as the training data (i.e., to define the trophic-niche-space models), (2) assumed all unobserved links between pairs of species were absent links, (3) ran the analysis only once on the full Serengeti dataset, and (4) prohibited impossible links based on each species' diet — we only treated species as predators if they preyed on vertebrates; we did not allow impossible links from vertebrates to strictly herbivorous or insectivorous species (e.g., water buffalo are strictly herbivorous, and so we did not assign vertebrate prey to this species).

In both the *GloBI* and Serengeti validations, the new model performed better than the simple body-size model (i.e., true skill statistics for the new model and the simple model were 0.38 vs 0.37, and 0.60 vs 0.58 for the *GloBI* and Serengeti data, respectively; Supplementary Table S5). Thus, we adopted the new model that incorporates body size and predator class to infer potential trophic links.

Figure S1 flowchart for building and analysing Naracoorte network models

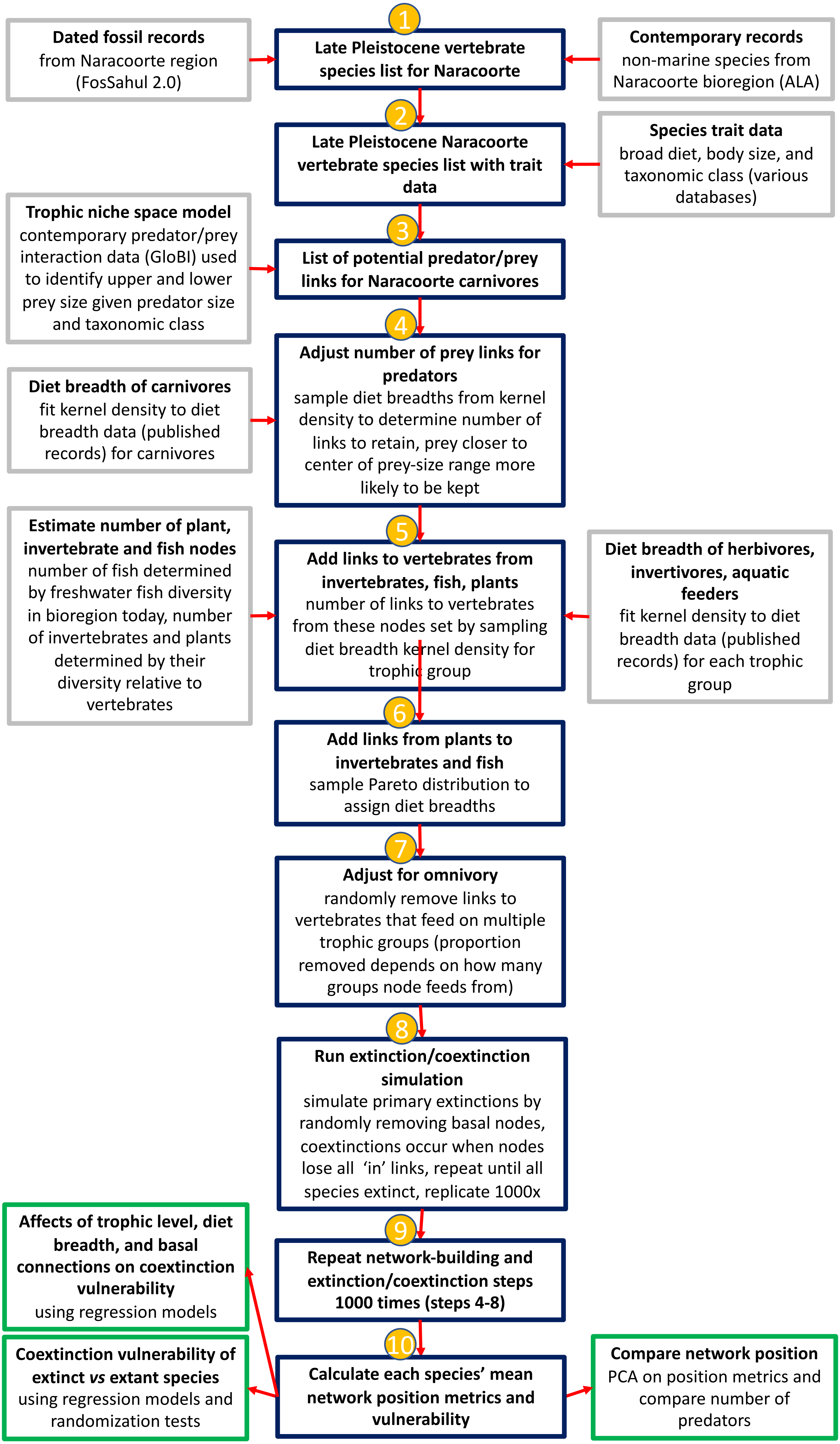

Figure S2 relationship between predator and prey body mass for predators from different taxonomic classes

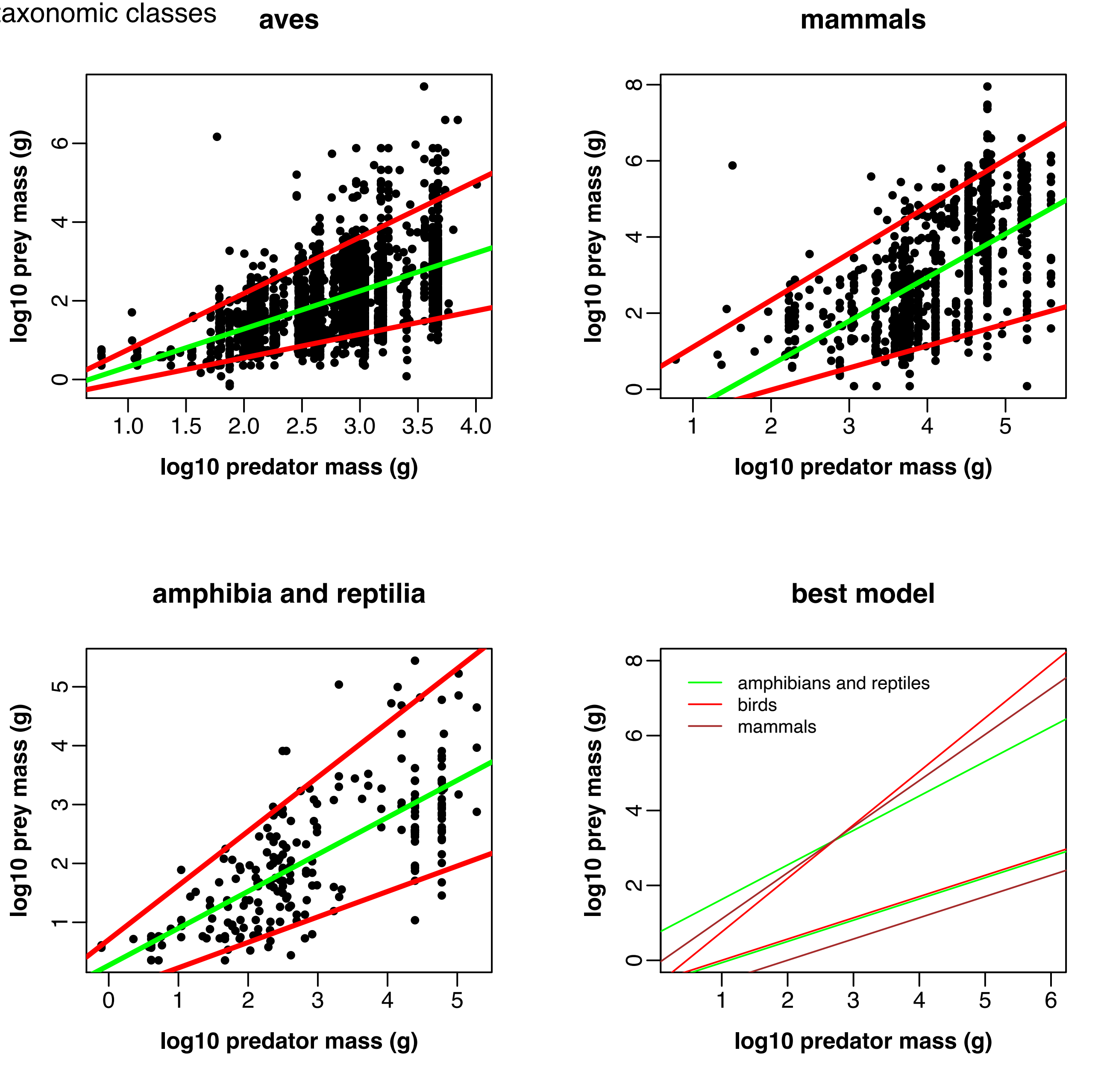

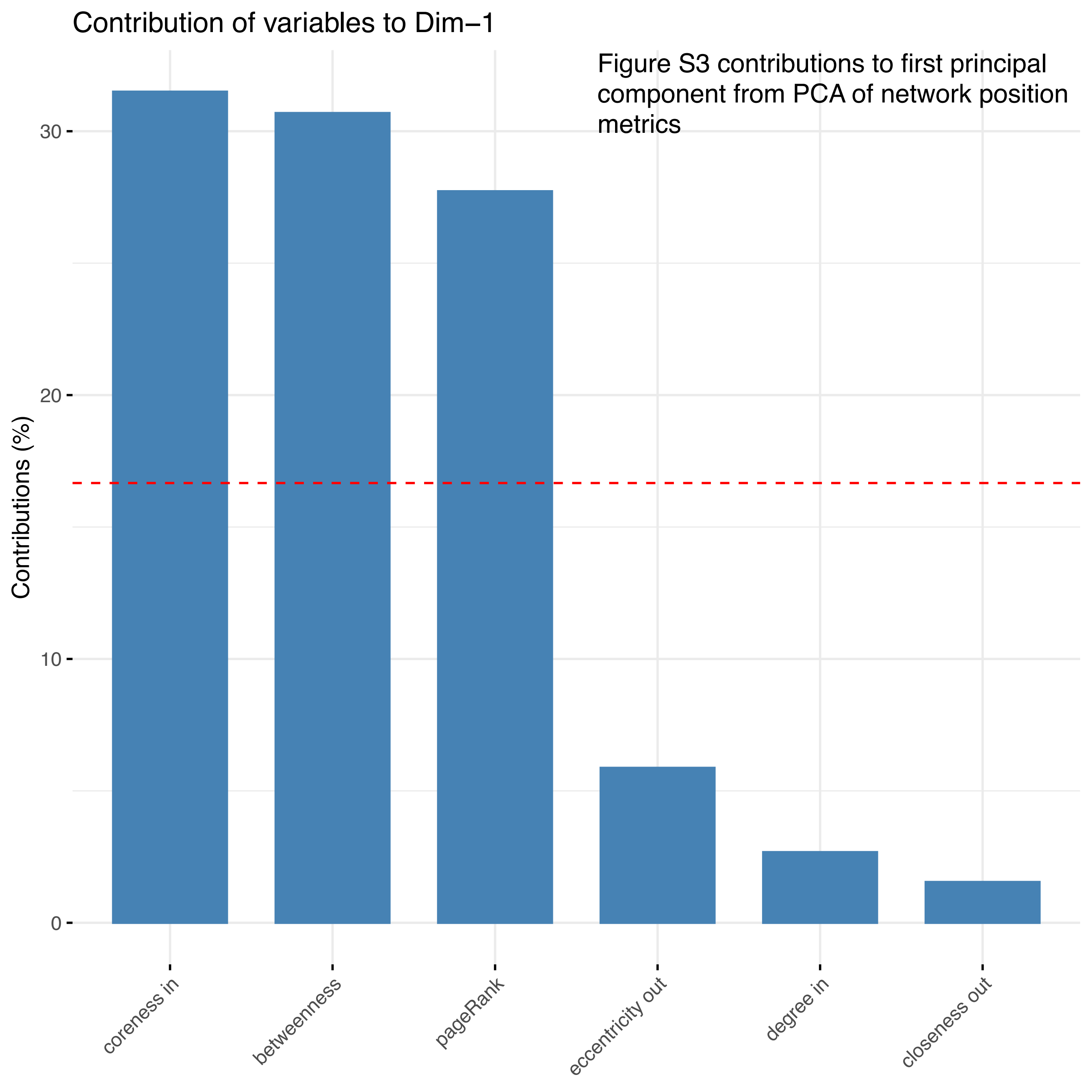

Figure S3 contributions to first principal component from PCA of network position metrics

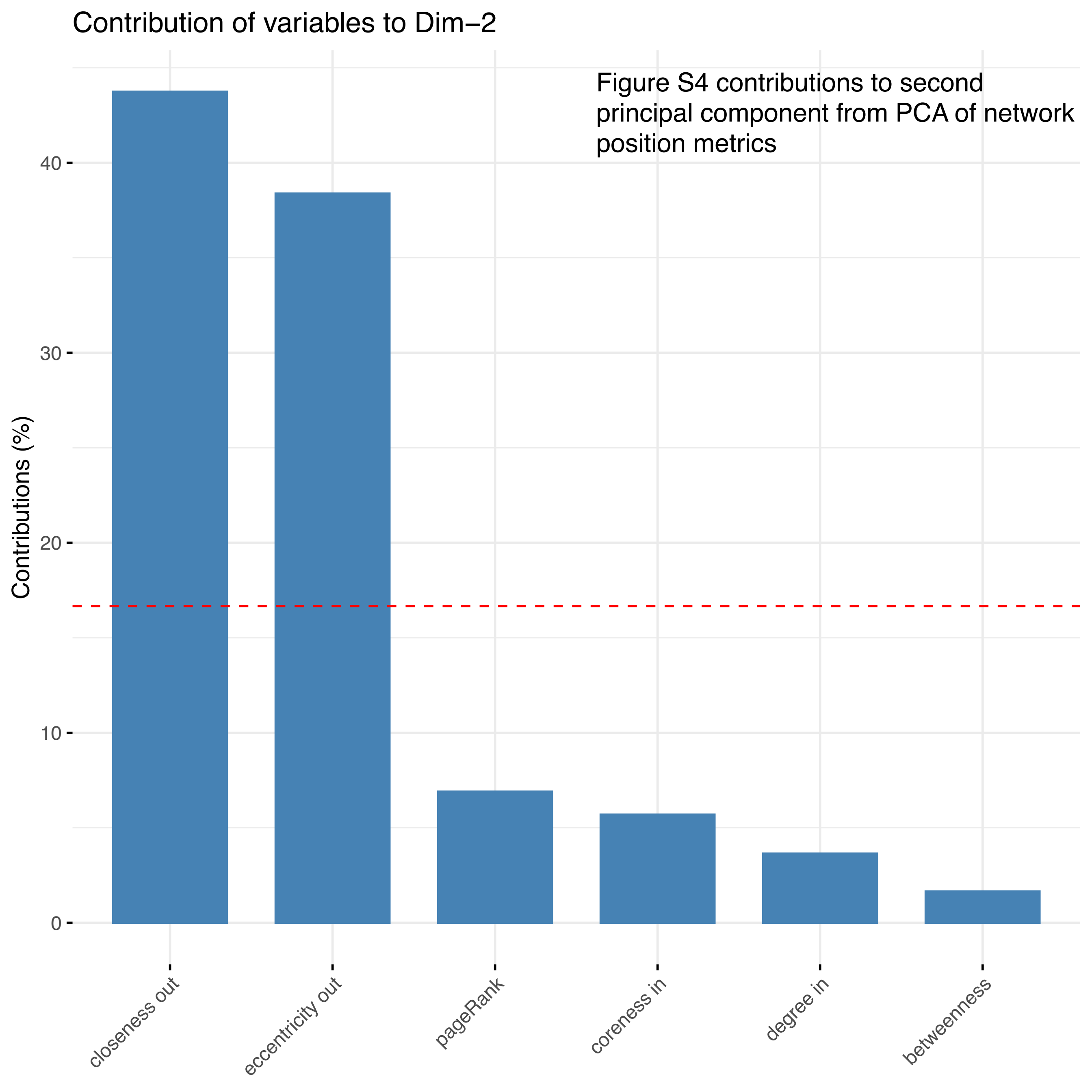

Figure S4 contributions to second principal component from PCA of network position metrics

**Supplementary figure S5: Temperature and precipitation anomalies in the Naracoorte region over the last 120,000 years**

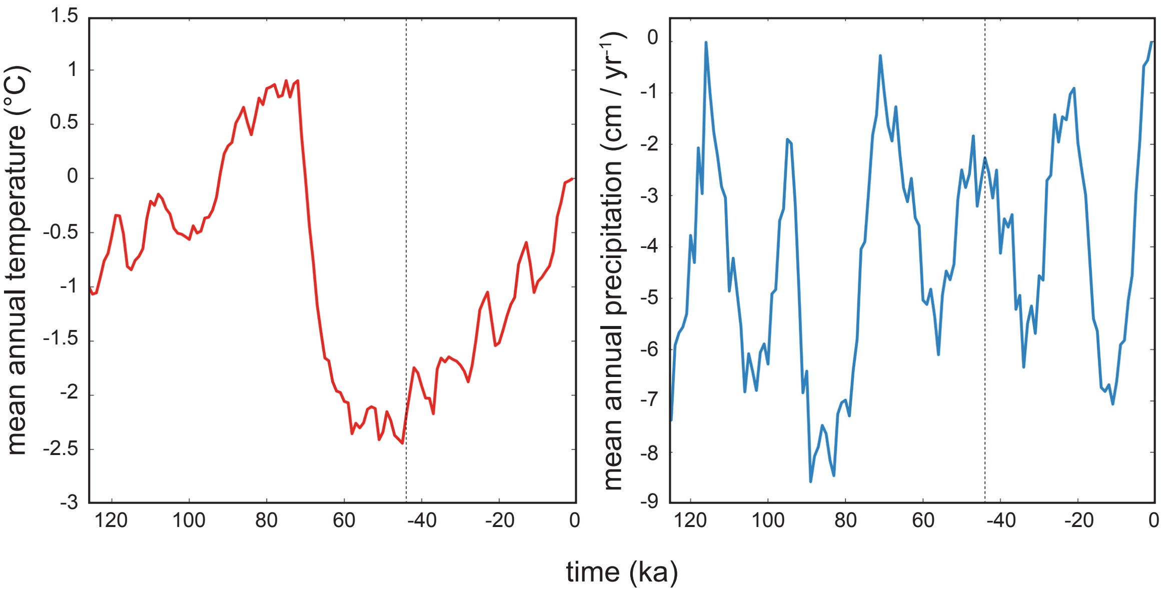

Naracoorte temperature and precipitation anomalies (relative to 1000 years ago) over the last 120,000 years. We hindcasted these climate variables for the Naracoorte region using a transient LOVECLIM Earth-system model[1,2]. The dotted vertical lines indicate the estimated date of megafauna extinction in this region (~ 44,000 years ago). Temperature reached a minimum immediately before the megafauna extinct (left panel), whereas the amount and rate of change in precipitation were not extreme at this time compared to earlier or later (right panel).

[1]Goosse, H. *et al.* Description of the Earth system model of intermediate complexity LOVECLIM version 1.2. *Geosci. Model Dev.* **3**, 603–633 (2010).

[2]Saltré, F. *et al.* Climate-human interaction associated with southeast Australian megafauna extinction patterns. *Nat. Commun.* **10**, 5311 (2019).

Table S1 Pre-extinction assemblage

| SciName | Class | Status (Late Pleistocene) | mean.mass.kg | eat.plant | eat.invert | eat.vert | eat.FishInv |
|---|---|---|---|---|---|---|---|
| Acritoscincus | Reptilia | Extant | 0.0029 | FALSE | TRUE | FALSE | FALSE |
| Acrobates py | Mammalia | Extant | 0.012 | TRUE | TRUE | FALSE | FALSE |
| Amphiboluru | Reptilia | Extant | 0.034711505 | FALSE | TRUE | TRUE | FALSE |
| Amphiboluru | Reptilia | Extant | 0.030502774 | TRUE | TRUE | TRUE | FALSE |
| Anilios bicolo | Reptilia | Extant | 0.00401829 | FALSE | TRUE | FALSE | FALSE |
| Anilios bitub | Reptilia | Extant | 0.008256638 | FALSE | TRUE | FALSE | FALSE |
| Antechinus a | Mammalia | Extant | 0.018 | TRUE | TRUE | FALSE | FALSE |
| Antechinus f | Mammalia | Extant | 0.04433 | TRUE | TRUE | TRUE | FALSE |
| Antechinus n | Mammalia | Extant | 0.0533 | FALSE | TRUE | TRUE | FALSE |
| Antechinus s | Mammalia | Extant | 0.053 | TRUE | TRUE | TRUE | FALSE |
| Aprasia strio | Reptilia | Extant | 0.007231299 | FALSE | TRUE | TRUE | FALSE |
| Austrelaps su | Reptilia | Extant | 0.63252 | FALSE | FALSE | TRUE | FALSE |
| Caligavis chr | Aves | Extant | 0.0162 | TRUE | TRUE | FALSE | FALSE |
| Cercartetus c | Mammalia | Extant | 0.013 | TRUE | TRUE | TRUE | FALSE |
| Cercartetus l | Mammalia | Extant | 0.0085 | TRUE | TRUE | TRUE | FALSE |
| Cercartetus n | Mammalia | Extant | 0.04 | TRUE | TRUE | TRUE | FALSE |
| Chalinolobus | Mammalia | Extant | 0.00891 | FALSE | TRUE | FALSE | FALSE |
| Chelodina lo | Reptilia | Extant | 0.602 | FALSE | TRUE | TRUE | FALSE |
| Christinus m | Reptilia | Extant | 0.00368 | FALSE | TRUE | FALSE | FALSE |
| Chrysococcyx | Aves | Extant | 0.023 | FALSE | TRUE | FALSE | FALSE |
| Chrysococcyx | Aves | Extant | 0.0242 | FALSE | TRUE | FALSE | FALSE |
| Chrysococcyx | Aves | Extant | 0.0308 | FALSE | TRUE | FALSE | FALSE |
| Cinclosoma c | Aves | Extant | 0.074515278 | TRUE | TRUE | FALSE | FALSE |
| Conilurus alb | Mammalia | Extant | 0.2 | TRUE | TRUE | FALSE | FALSE |
| Coturnix ypsi | Aves | Extant | 0.105 | TRUE | TRUE | FALSE | FALSE |
| Crinia parins | Amphibia | Extant | 7.00E-04 | FALSE | TRUE | FALSE | FALSE |
| Crinia signife | Amphibia | Extant | 0.00071 | FALSE | TRUE | FALSE | FALSE |
| Ctenophorus | Reptilia | Extant | 0.00409 | FALSE | TRUE | FALSE | FALSE |

| | | | | | | | |
|---|---|---|---|---|---|---|---|
| Ctenophorus | Reptilia | Extant | 0.008 | FALSE | TRUE | TRUE | FALSE |
| Ctenotus orie | Reptilia | Extant | 0.009461111 | FALSE | TRUE | TRUE | FALSE |
| Ctenotus rob | Reptilia | Extant | 0.032308931 | TRUE | TRUE | TRUE | FALSE |
| Ctenotus spa | Reptilia | Extant | 0.017258379 | FALSE | TRUE | TRUE | FALSE |
| Ctenotus tae | Reptilia | Extant | 0.002980311 | FALSE | TRUE | FALSE | FALSE |
| Ctenotus ube | Reptilia | Extant | 0.01 | FALSE | TRUE | TRUE | FALSE |
| Dasyurus ma | Mammalia | Extant | 2.65 | TRUE | TRUE | TRUE | FALSE |
| Dasyurus viv | Mammalia | Extant | 1.13 | TRUE | TRUE | TRUE | FALSE |
| Delma austra | Reptilia | Extant | 0.004235315 | FALSE | TRUE | FALSE | FALSE |
| Delma impar | Reptilia | Extant | 0.007 | FALSE | TRUE | TRUE | FALSE |
| Delma inorna | Reptilia | Extant | 0.007461061 | FALSE | TRUE | TRUE | FALSE |
| Drysdalia cor | Reptilia | Extant | 0.036921771 | FALSE | FALSE | TRUE | FALSE |
| Drysdalia ma | Reptilia | Extant | 0.017501008 | FALSE | FALSE | TRUE | FALSE |
| Emydura ma | Reptilia | Extant | 3.2 | TRUE | TRUE | FALSE | FALSE |
| Falsistrellus | Mammalia | Extant | 0.02254 | FALSE | TRUE | FALSE | FALSE |
| Gallirallus pl | Aves | Extant | 0.175 | TRUE | TRUE | TRUE | TRUE |
| Gavicalis vire | Aves | Extant | 0.0243 | TRUE | TRUE | FALSE | FALSE |
| Geocrinia lae | Amphibia | Extant | 0.00092 | FALSE | TRUE | FALSE | FALSE |
| Gliciphila me | Aves | Extant | 0.0185 | TRUE | TRUE | FALSE | FALSE |
| Hemiergis de | Reptilia | Extant | 0.00161 | FALSE | TRUE | FALSE | FALSE |
| Hemiergis pe | Reptilia | Extant | 0.00375 | FALSE | TRUE | FALSE | FALSE |
| Himantopus | Aves | Extant | 0.178 | FALSE | TRUE | FALSE | FALSE |
| Hydromys ch | Mammalia | Extant | 0.85 | FALSE | TRUE | TRUE | TRUE |
| Hylacola cau | Aves | Extant | 0.0146875 | FALSE | TRUE | FALSE | FALSE |
| Hylacola pyrr | Aves | Extant | 0.017011111 | TRUE | TRUE | FALSE | FALSE |
| Isoodon obes | Mammalia | Extant | 1.25 | TRUE | TRUE | FALSE | FALSE |
| Lampropholi | Reptilia | Extant | 0.00116 | FALSE | TRUE | FALSE | FALSE |
| Lampropholi | Reptilia | Extant | 0.00106 | FALSE | TRUE | FALSE | FALSE |
| Lerista boug | Reptilia | Extant | 0.00157 | FALSE | TRUE | FALSE | FALSE |
| Limnodynast | Amphibia | Extant | 0.046714977 | FALSE | TRUE | TRUE | FALSE |

| | | | | | | | |
|---|---|---|---|---|---|---|---|
| Limnodynast | Amphibia | Extant | 0.0155 | FALSE | TRUE | TRUE | FALSE |
| Litoria ewing | Amphibia | Extant | 0.00169 | FALSE | TRUE | FALSE | FALSE |
| Litoria peron | Amphibia | Extant | 0.00791 | FALSE | TRUE | TRUE | FALSE |
| Litoria ranifo | Amphibia | Extant | 0.0742886 | FALSE | TRUE | TRUE | FALSE |
| Macropus gr | Mammalia | Extant | 10 | TRUE | FALSE | FALSE | FALSE |
| Mastacomys | Mammalia | Extant | 0.122 | TRUE | TRUE | FALSE | FALSE |
| Menetia grey | Reptilia | Extant | 0.000415 | FALSE | TRUE | FALSE | FALSE |
| Miniopterus | Mammalia | Extant | 0.0109 | FALSE | TRUE | FALSE | FALSE |
| Morethia ade | Reptilia | Extant | 0.0015 | FALSE | TRUE | FALSE | FALSE |
| Morethia bo | Reptilia | Extant | 0.0015 | FALSE | TRUE | FALSE | FALSE |
| Morethia obs | Reptilia | Extant | 0.002980311 | FALSE | TRUE | FALSE | FALSE |
| Mormopteru | Mammalia | Extant | 0.0088 | FALSE | TRUE | FALSE | FALSE |
| Neobatrachu | Amphibia | Extant | 0.007201685 | FALSE | TRUE | TRUE | FALSE |
| Neobatrachu | Amphibia | Extant | 0.007201685 | FALSE | TRUE | TRUE | FALSE |
| Nesoptilotis | Aves | Extant | 0.0216 | TRUE | TRUE | FALSE | FALSE |
| Notamacrop | Mammalia | Extant | 5.28 | TRUE | FALSE | FALSE | FALSE |
| Notamacrop | Mammalia | Extant | 16.8 | TRUE | FALSE | FALSE | FALSE |
| Notechis scu | Reptilia | Extant | 0.8355 | FALSE | FALSE | TRUE | FALSE |
| Notomys mit | Mammalia | Extant | 0.047 | TRUE | FALSE | FALSE | FALSE |
| Nyctophilus g | Mammalia | Extant | 0.008 | FALSE | TRUE | FALSE | FALSE |
| Nyctophilus g | Mammalia | Extant | 0.01132 | FALSE | TRUE | FALSE | FALSE |
| Ornithorhync | Mammalia | Extant | 1.48 | FALSE | TRUE | FALSE | TRUE |
| Parasuta flag | Reptilia | Extant | 0.012882496 | FALSE | FALSE | TRUE | FALSE |
| Parasuta nig | Reptilia | Extant | 0.022387211 | FALSE | FALSE | TRUE | FALSE |
| Parvipsitta p | Aves | Extant | 0.0449 | TRUE | FALSE | FALSE | FALSE |
| Parvipsitta p | Aves | Extant | 0.0395 | TRUE | FALSE | FALSE | FALSE |
| Perameles b | Mammalia | Extant | 0.226 | TRUE | TRUE | TRUE | FALSE |
| Perameles g | Mammalia | Extant | 0.75 | TRUE | TRUE | TRUE | FALSE |
| Petaurus aus | Mammalia | Extant | 0.571 | TRUE | TRUE | FALSE | FALSE |
| Petaurus bre | Mammalia | Extant | 0.11 | TRUE | TRUE | FALSE | FALSE |

| | | | | | | | |
|---|---|---|---|---|---|---|---|
| Petroica boo | Aves | Extant | 0.0132 | FALSE | TRUE | FALSE | FALSE |
| Phalacrocora | Aves | Extant | 0.688 | FALSE | TRUE | TRUE | TRUE |
| Phascogale c | Mammalia | Extant | 0.0433 | TRUE | TRUE | TRUE | FALSE |
| Phascogale t | Mammalia | Extant | 0.19 | TRUE | TRUE | TRUE | FALSE |
| Phascolarcto | Mammalia | Extant | 8.45 | TRUE | FALSE | FALSE | FALSE |
| Phylidonyris | Aves | Extant | 0.015011304 | TRUE | TRUE | FALSE | FALSE |
| Pogona barb | Reptilia | Extant | 0.29 | TRUE | TRUE | TRUE | FALSE |
| Pogona vittic | Reptilia | Extant | 0.359080138 | TRUE | TRUE | TRUE | FALSE |
| Potorous pla | Mammalia | Extant | 0.5 | TRUE | TRUE | FALSE | FALSE |
| Potorous trid | Mammalia | Extant | 1.05 | TRUE | TRUE | FALSE | FALSE |
| Pseudemoia | Reptilia | Extant | 0.004 | FALSE | TRUE | FALSE | FALSE |
| Pseudemoia | Reptilia | Extant | 0.004056528 | FALSE | TRUE | FALSE | FALSE |
| Pseudocheiru | Mammalia | Extant | 0.8 | TRUE | FALSE | FALSE | FALSE |
| Pseudomys a | Mammalia | Extant | 0.02 | TRUE | TRUE | FALSE | FALSE |
| Pseudomys a | Mammalia | Extant | 0.053 | TRUE | TRUE | FALSE | FALSE |
| Pseudomys a | Mammalia | Extant | 0.053 | TRUE | TRUE | FALSE | FALSE |
| Pseudomys f | Mammalia | Extant | 0.06875 | TRUE | TRUE | FALSE | FALSE |
| Pseudomys g | Mammalia | Extant | 0.05 | TRUE | TRUE | FALSE | FALSE |
| Pseudomys s | Mammalia | Extant | 0.064 | TRUE | TRUE | FALSE | FALSE |
| Pseudonaja t | Reptilia | Extant | 0.469 | FALSE | FALSE | TRUE | FALSE |
| Pseudophryn | Amphibia | Extant | 0.002 | FALSE | TRUE | FALSE | FALSE |
| Pseudophryn | Amphibia | Extant | 0.003541306 | FALSE | TRUE | FALSE | FALSE |
| Pteropus pol | Mammalia | Extant | 0.702 | TRUE | FALSE | FALSE | FALSE |
| Ptilotula fusc | Aves | Extant | 0.016249575 | TRUE | TRUE | FALSE | FALSE |
| Ptilotula orn | Aves | Extant | 0.017193188 | TRUE | TRUE | FALSE | FALSE |
| Ptilotula pen | Aves | Extant | 0.0183 | TRUE | TRUE | FALSE | FALSE |
| Pygopus lepi | Reptilia | Extant | 0.020098147 | FALSE | TRUE | TRUE | FALSE |
| Rattus fuscip | Mammalia | Extant | 0.133 | TRUE | TRUE | FALSE | FALSE |
| Rattus leuco | Mammalia | Extant | 0.201 | TRUE | TRUE | FALSE | FALSE |
| Rattus lutreo | Mammalia | Extant | 0.125 | TRUE | TRUE | FALSE | FALSE |

| | | | | | | | |
|---|---|---|---|---|---|---|---|
| Rattus tunne | Mammalia | Extant | 0.124 | TRUE | TRUE | FALSE | FALSE |
| Rhipidura alb | Aves | Extant | 0.0078 | FALSE | TRUE | FALSE | FALSE |
| Saccolaimus | Mammalia | Extant | 0.0452 | FALSE | TRUE | FALSE | FALSE |
| Scotorepens | Mammalia | Extant | 0.0118 | FALSE | TRUE | FALSE | FALSE |
| Sminthopsis | Mammalia | Extant | 0.016 | FALSE | TRUE | TRUE | FALSE |
| Sminthopsis | Mammalia | Extant | 0.02336 | FALSE | TRUE | TRUE | FALSE |
| Sminthopsis | Mammalia | Extant | 0.024 | FALSE | TRUE | TRUE | FALSE |
| Sminthopsis | Mammalia | Extant | 0.017 | FALSE | TRUE | TRUE | FALSE |
| Tachyglossus | Mammalia | Extant | 4.5 | FALSE | TRUE | FALSE | FALSE |
| Tadarida aus | Mammalia | Extant | 0.0364 | FALSE | TRUE | FALSE | FALSE |
| Threskiornis | Aves | Extant | 1.81 | FALSE | TRUE | FALSE | TRUE |
| Thylacinus cy | Mammalia | Extant | 30 | FALSE | FALSE | TRUE | FALSE |
| Tiliqua nigro | Reptilia | Extant | 0.481061145 | TRUE | TRUE | TRUE | FALSE |
| Tiliqua occip | Reptilia | Extant | 0.584923789 | TRUE | TRUE | TRUE | FALSE |
| Tiliqua rugos | Reptilia | Extant | 0.61695 | TRUE | TRUE | TRUE | FALSE |
| Tiliqua scinc | Reptilia | Extant | 0.499833333 | TRUE | TRUE | TRUE | FALSE |
| Trichosurus v | Mammalia | Extant | 2.32 | TRUE | FALSE | FALSE | FALSE |
| Tympanocryp | Reptilia | Extant | 0.00972 | FALSE | TRUE | TRUE | FALSE |
| Underwoodis | Reptilia | Extant | 0.00961 | FALSE | TRUE | TRUE | FALSE |
| Varanus gou | Reptilia | Extant | 0.82106 | FALSE | TRUE | TRUE | FALSE |
| Varanus rose | Reptilia | Extant | 1.109 | FALSE | TRUE | TRUE | FALSE |
| Varanus vari | Reptilia | Extant | 6.343 | FALSE | TRUE | TRUE | FALSE |
| Vespadelus d | Mammalia | Extant | 0.00606 | FALSE | TRUE | FALSE | FALSE |
| Vespadelus r | Mammalia | Extant | 0.00505 | FALSE | TRUE | FALSE | FALSE |
| Vespadelus v | Mammalia | Extant | 0.00377 | FALSE | TRUE | FALSE | FALSE |
| Vombatus ur | Mammalia | Extant | 26 | TRUE | FALSE | FALSE | FALSE |
| Wallabia bic | Mammalia | Extant | 14.63 | TRUE | FALSE | FALSE | FALSE |
| Acanthageny | Aves | Extant | 0.044730602 | TRUE | TRUE | FALSE | FALSE |
| Acanthiza ap | Aves | Extant | 0.0072 | TRUE | TRUE | FALSE | FALSE |
| Acanthiza ch | Aves | Extant | 0.0091 | TRUE | TRUE | FALSE | FALSE |

| | | | | | | | |
|---|---|---|---|---|---|---|---|
| Acanthiza ire | Aves | Extant | 0.0061 | FALSE | TRUE | FALSE | FALSE |
| Acanthiza lin | Aves | Extant | 0.0073 | TRUE | TRUE | FALSE | FALSE |
| Acanthiza na | Aves | Extant | 0.0065 | TRUE | TRUE | FALSE | FALSE |
| Acanthiza pu | Aves | Extant | 0.009 | TRUE | TRUE | FALSE | FALSE |
| Acanthiza re | Aves | Extant | 0.0074 | TRUE | TRUE | FALSE | FALSE |
| Acanthiza ur | Aves | Extant | 0.006425 | TRUE | TRUE | FALSE | FALSE |
| Acanthorhyn | Aves | Extant | 0.0105 | TRUE | TRUE | FALSE | FALSE |
| Accipiter cirr | Aves | Extant | 0.18 | FALSE | TRUE | TRUE | FALSE |
| Accipiter fas | Aves | Extant | 0.371 | FALSE | FALSE | TRUE | FALSE |
| Accipiter nov | Aves | Extant | 0.565 | FALSE | FALSE | TRUE | FALSE |
| Acrocephalus | Aves | Extant | 0.0189 | FALSE | TRUE | FALSE | FALSE |
| Actitis hypol | Aves | Extant | 0.0417163 | FALSE | TRUE | FALSE | TRUE |
| Aegotheles c | Aves | Extant | 0.042833333 | FALSE | TRUE | FALSE | FALSE |
| Amytornis st | Aves | Extant | 0.01855 | TRUE | TRUE | FALSE | FALSE |
| Anas castane | Aves | Extant | 0.638 | TRUE | TRUE | FALSE | TRUE |
| Anas gracilis | Aves | Extant | 0.493 | TRUE | TRUE | FALSE | TRUE |
| Anas rhyncho | Aves | Extant | 0.646 | TRUE | TRUE | FALSE | TRUE |
| Anas superci | Aves | Extant | 1.06 | TRUE | TRUE | FALSE | TRUE |
| Anhinga nova | Aves | Extant | 1.54 | FALSE | FALSE | FALSE | TRUE |
| Anseranas se | Aves | Extant | 2.42 | TRUE | FALSE | FALSE | FALSE |
| Anthochaera | Aves | Extant | 0.090904012 | TRUE | TRUE | FALSE | FALSE |
| Anthochaera | Aves | Extant | 0.0697 | TRUE | TRUE | FALSE | FALSE |
| Anthochaera | Aves | Extant | 0.0619 | TRUE | TRUE | FALSE | FALSE |
| Anthus novae | Aves | Extant | 0.0257 | TRUE | TRUE | FALSE | FALSE |
| Aphelocepha | Aves | Extant | 0.01235 | TRUE | TRUE | FALSE | FALSE |
| Apus pacificu | Aves | Extant | 0.0435 | FALSE | TRUE | FALSE | FALSE |
| Aquila audax | Aves | Extant | 3.63 | FALSE | FALSE | TRUE | FALSE |
| Ardea alba | Aves | Extant | 0.93 | FALSE | TRUE | TRUE | TRUE |
| Ardea interm | Aves | Extant | 0.45 | FALSE | TRUE | FALSE | TRUE |
| Ardea pacific | Aves | Extant | 0.881 | FALSE | TRUE | TRUE | TRUE |

| | | | | | | | |
|---|---|---|---|---|---|---|---|
| Ardeotis aus | Aves | Extant | 4.89 | TRUE | TRUE | TRUE | FALSE |
| Arenaria inte | Aves | Extant | 0.119 | FALSE | TRUE | FALSE | FALSE |
| Artamus cine | Aves | Extant | 0.03533475 | TRUE | TRUE | FALSE | FALSE |
| Artamus cya | Aves | Extant | 0.034 | TRUE | TRUE | FALSE | FALSE |
| Artamus leu | Aves | Extant | 0.0428 | TRUE | TRUE | FALSE | FALSE |
| Artamus per | Aves | Extant | 0.0347 | TRUE | TRUE | FALSE | FALSE |
| Artamus sup | Aves | Extant | 0.0353 | TRUE | TRUE | FALSE | FALSE |
| Aythya austr | Aves | Extant | 0.87 | TRUE | TRUE | FALSE | TRUE |
| Barnardius z | Aves | Extant | 0.151 | TRUE | TRUE | FALSE | FALSE |
| Biziura lobat | Aves | Extant | 1.67 | TRUE | FALSE | FALSE | TRUE |
| Botaurus poi | Aves | Extant | 1.11 | FALSE | FALSE | FALSE | TRUE |
| Burhinus gra | Aves | Extant | 0.673 | TRUE | TRUE | FALSE | FALSE |
| Cacatua gale | Aves | Extant | 0.735 | TRUE | FALSE | FALSE | FALSE |
| Cacatua sang | Aves | Extant | 0.497 | TRUE | FALSE | FALSE | FALSE |
| Cacatua tenu | Aves | Extant | 0.567 | TRUE | FALSE | FALSE | FALSE |
| Cacomantis f | Aves | Extant | 0.0499 | FALSE | TRUE | FALSE | FALSE |
| Cacomantis p | Aves | Extant | 0.0876 | FALSE | TRUE | FALSE | FALSE |
| Calamanthus | Aves | Extant | 0.014472619 | FALSE | TRUE | FALSE | FALSE |
| Calamanthus | Aves | Extant | 0.019125 | FALSE | TRUE | FALSE | FALSE |
| Calidris acum | Aves | Extant | 0.0671 | TRUE | TRUE | FALSE | TRUE |
| Calidris ferru | Aves | Extant | 0.06229732 | TRUE | TRUE | FALSE | TRUE |
| Calidris mela | Aves | Extant | 0.071785266 | TRUE | TRUE | FALSE | TRUE |
| Calidris rufic | Aves | Extant | 0.028495478 | TRUE | TRUE | FALSE | TRUE |
| Calidris subm | Aves | Extant | 0.024638889 | TRUE | FALSE | FALSE | TRUE |
| Callocephalo | Aves | Extant | 0.257 | TRUE | FALSE | FALSE | FALSE |
| Calyptorhync | Aves | Extant | 0.682 | TRUE | FALSE | FALSE | FALSE |
| Calyptorhync | Aves | Extant | 0.661 | TRUE | TRUE | FALSE | FALSE |
| Cereopsis no | Aves | Extant | 5.36 | TRUE | FALSE | FALSE | FALSE |
| Certhionyx va | Aves | Extant | 0.026383182 | TRUE | TRUE | FALSE | FALSE |
| Ceyx azureus | Aves | Extant | 0.035202333 | FALSE | FALSE | FALSE | TRUE |

| | | | | | | | |
|---|---|---|---|---|---|---|---|
| Charadrius b | Aves | Extant | 0.070839306 | FALSE | TRUE | FALSE | TRUE |
| Charadrius ru | Aves | Extant | 0.037451045 | FALSE | TRUE | FALSE | TRUE |
| Charadrius ve | Aves | Extant | 0.095 | FALSE | TRUE | FALSE | FALSE |
| Chenonetta j | Aves | Extant | 0.808 | TRUE | FALSE | FALSE | FALSE |
| Cheramoeca | Aves | Extant | 0.014018254 | FALSE | TRUE | FALSE | FALSE |
| Chlidonias hy | Aves | Extant | 0.0856 | FALSE | TRUE | TRUE | TRUE |
| Chlidonias le | Aves | Extant | 0.0649 | FALSE | TRUE | TRUE | TRUE |
| Chroicocepha | Aves | Extant | 0.287 | FALSE | TRUE | TRUE | TRUE |
| Cincloramph | Aves | Extant | 0.0532 | TRUE | TRUE | FALSE | FALSE |
| Cincloramph | Aves | Extant | 0.0297 | FALSE | TRUE | FALSE | FALSE |
| Circus approx | Aves | Extant | 0.754 | FALSE | FALSE | TRUE | FALSE |
| Circus assim | Aves | Extant | 0.568 | FALSE | FALSE | TRUE | FALSE |
| Cisticola exil | Aves | Extant | 0.0078 | TRUE | TRUE | FALSE | FALSE |
| Cladorhynchu | Aves | Extant | 0.317 | FALSE | FALSE | FALSE | TRUE |
| Climacteris a | Aves | Extant | 0.020825 | FALSE | TRUE | FALSE | FALSE |
| Climacteris p | Aves | Extant | 0.032 | TRUE | TRUE | FALSE | FALSE |
| Colluricincla | Aves | Extant | 0.0675 | TRUE | TRUE | TRUE | FALSE |
| Coracina nov | Aves | Extant | 0.115 | TRUE | TRUE | FALSE | FALSE |
| Coracina pap | Aves | Extant | 0.0659 | TRUE | TRUE | FALSE | FALSE |
| Corcorax me | Aves | Extant | 0.361 | TRUE | TRUE | FALSE | FALSE |
| Cormobates | Aves | Extant | 0.02 | TRUE | TRUE | FALSE | FALSE |
| Corvus benne | Aves | Extant | 0.396 | TRUE | TRUE | TRUE | FALSE |
| Corvus coron | Aves | Extant | 0.593 | TRUE | TRUE | TRUE | FALSE |
| Corvus mello | Aves | Extant | 0.534 | FALSE | TRUE | TRUE | FALSE |
| Corvus tasma | Aves | Extant | 0.675 | TRUE | TRUE | TRUE | FALSE |
| Coturnix pect | Aves | Extant | 0.101 | TRUE | TRUE | FALSE | FALSE |
| Cracticus tibi | Aves | Extant | 0.28 | TRUE | TRUE | TRUE | FALSE |
| Cracticus tor | Aves | Extant | 0.090785948 | FALSE | TRUE | TRUE | FALSE |
| Cygnus atrat | Aves | Extant | 5.66 | TRUE | FALSE | FALSE | FALSE |
| Dacelo novae | Aves | Extant | 0.312 | FALSE | TRUE | TRUE | TRUE |

| | | | | | | | |
|---|---|---|---|---|---|---|---|
| Daphoenositt | Aves | Extant | 0.0121 | FALSE | TRUE | FALSE | FALSE |
| Dendrocygna | Aves | Extant | 0.733 | TRUE | FALSE | FALSE | TRUE |
| Dicaeum hiru | Aves | Extant | 0.0088 | TRUE | TRUE | FALSE | FALSE |
| Dromaius no | Aves | Extant | 35.5 | TRUE | TRUE | FALSE | FALSE |
| Drymodes br | Aves | Extant | 0.033476316 | TRUE | TRUE | FALSE | FALSE |
| Egretta garz | Aves | Extant | 0.482 | FALSE | TRUE | FALSE | TRUE |
| Egretta nova | Aves | Extant | 0.571 | FALSE | TRUE | FALSE | TRUE |
| Elanus axilla | Aves | Extant | 0.275 | FALSE | TRUE | TRUE | FALSE |
| Elanus script | Aves | Extant | 0.316 | FALSE | TRUE | TRUE | FALSE |
| Elseyornis m | Aves | Extant | 0.032500935 | FALSE | TRUE | FALSE | TRUE |
| Entomyzon c | Aves | Extant | 0.103 | TRUE | TRUE | FALSE | FALSE |
| Eolophus ros | Aves | Extant | 0.306 | TRUE | FALSE | FALSE | FALSE |
| Eopsaltria au | Aves | Extant | 0.0194 | FALSE | TRUE | FALSE | FALSE |
| Epthianura a | Aves | Extant | 0.0133 | FALSE | TRUE | FALSE | FALSE |
| Epthianura a | Aves | Extant | 0.010514018 | FALSE | TRUE | FALSE | FALSE |
| Epthianura ti | Aves | Extant | 0.010711667 | TRUE | TRUE | FALSE | FALSE |
| Erythrogonys | Aves | Extant | 0.050404407 | FALSE | FALSE | FALSE | TRUE |
| Eurostopodu | Aves | Extant | 0.093 | FALSE | TRUE | FALSE | FALSE |
| Falco berigor | Aves | Extant | 0.574 | FALSE | TRUE | TRUE | FALSE |
| Falco cenchr | Aves | Extant | 0.179 | FALSE | TRUE | TRUE | FALSE |
| Falco hypole | Aves | Extant | 0.466 | FALSE | FALSE | TRUE | FALSE |
| Falco longipe | Aves | Extant | 0.244 | FALSE | TRUE | TRUE | FALSE |
| Falco peregr | Aves | Extant | 1.09 | FALSE | FALSE | TRUE | FALSE |
| Falco subnig | Aves | Extant | 0.738 | FALSE | FALSE | TRUE | FALSE |
| Falcunculus f | Aves | Extant | 0.025651121 | TRUE | TRUE | FALSE | FALSE |
| Fulica atra | Aves | Extant | 0.651 | TRUE | FALSE | FALSE | FALSE |
| Gallinago ha | Aves | Extant | 0.162 | TRUE | TRUE | FALSE | TRUE |
| Gallinula ten | Aves | Extant | 0.432 | TRUE | FALSE | FALSE | TRUE |
| Gelochelidor | Aves | Extant | 0.2 | FALSE | TRUE | FALSE | TRUE |
| Geopelia cur | Aves | Extant | 0.032108813 | TRUE | FALSE | FALSE | FALSE |

| | | | | | | | |
|---|---|---|---|---|---|---|---|
| Geopelia stri | Aves | Extant | 0.0454 | TRUE | FALSE | FALSE | FALSE |
| Gerygone oli | Aves | Extant | 0.0065 | FALSE | TRUE | FALSE | FALSE |
| Glareola ma | Aves | Extant | 0.075319835 | FALSE | TRUE | FALSE | FALSE |
| Glossopsitta | Aves | Extant | 0.070775 | TRUE | FALSE | FALSE | FALSE |
| Grallina cyar | Aves | Extant | 0.087982587 | FALSE | TRUE | FALSE | TRUE |
| Grantiella pi | Aves | Extant | 0.020666667 | TRUE | TRUE | FALSE | FALSE |
| Grus rubicun | Aves | Extant | 6.25 | TRUE | TRUE | TRUE | TRUE |
| Haliastur sph | Aves | Extant | 0.769 | FALSE | TRUE | TRUE | TRUE |
| Hieraaetus m | Aves | Extant | 0.832 | FALSE | TRUE | TRUE | FALSE |
| Hirundapus c | Aves | Extant | 0.119 | FALSE | TRUE | FALSE | FALSE |
| Hirundo neox | Aves | Extant | 0.01455 | FALSE | TRUE | FALSE | FALSE |
| Hydroprogne | Aves | Extant | 0.752 | FALSE | FALSE | FALSE | TRUE |
| Ixobrychus du | Aves | Extant | 0.084 | FALSE | FALSE | FALSE | TRUE |
| Lalage tricolo | Aves | Extant | 0.0255 | TRUE | TRUE | FALSE | FALSE |
| Lathamus dis | Aves | Extant | 0.067287319 | TRUE | FALSE | FALSE | FALSE |
| Leipoa ocella | Aves | Extant | 1.92 | TRUE | TRUE | FALSE | FALSE |
| Lewinia pect | Aves | Extant | 0.087833333 | FALSE | TRUE | FALSE | TRUE |
| Lichenostom | Aves | Extant | 0.019772256 | TRUE | TRUE | FALSE | FALSE |
| Lichenostom | Aves | Extant | 0.026826788 | TRUE | TRUE | FALSE | FALSE |
| Limosa lappo | Aves | Extant | 0.317 | FALSE | TRUE | FALSE | FALSE |
| Limosa limos | Aves | Extant | 0.263 | FALSE | TRUE | FALSE | TRUE |
| Lophoictinia | Aves | Extant | 0.67 | FALSE | TRUE | TRUE | FALSE |
| Malacorhync | Aves | Extant | 0.377 | TRUE | FALSE | FALSE | TRUE |
| Malurus cyar | Aves | Extant | 0.0106 | TRUE | TRUE | FALSE | FALSE |
| Malurus lam | Aves | Extant | 0.007984377 | FALSE | TRUE | FALSE | FALSE |
| Malurus sple | Aves | Extant | 0.009175312 | FALSE | TRUE | FALSE | FALSE |
| Manorina fla | Aves | Extant | 0.0573525 | TRUE | TRUE | FALSE | FALSE |
| Manorina me | Aves | Extant | 0.0713 | TRUE | TRUE | TRUE | FALSE |
| Megalurus g | Aves | Extant | 0.0126 | FALSE | TRUE | FALSE | FALSE |
| Melanodryas | Aves | Extant | 0.0193 | FALSE | TRUE | FALSE | FALSE |

| | | | | | | | |
|---|---|---|---|---|---|---|---|
| Melithreptus | Aves | Extant | 0.0153 | TRUE | TRUE | FALSE | FALSE |
| Melithreptus | Aves | Extant | 0.0196 | TRUE | TRUE | FALSE | FALSE |
| Melithreptus | Aves | Extant | 0.0149 | TRUE | TRUE | FALSE | FALSE |
| Melopsittacu | Aves | Extant | 0.028792403 | TRUE | FALSE | FALSE | FALSE |
| Merops orna | Aves | Extant | 0.028 | FALSE | TRUE | FALSE | FALSE |
| Microeca fas | Aves | Extant | 0.0145 | FALSE | TRUE | FALSE | FALSE |
| Milvus migra | Aves | Extant | 0.847 | FALSE | TRUE | TRUE | TRUE |
| Mirafra javar | Aves | Extant | 0.0218 | TRUE | TRUE | FALSE | FALSE |
| Myiagra cyar | Aves | Extant | 0.0174 | FALSE | TRUE | FALSE | FALSE |
| Myiagra inqu | Aves | Extant | 0.0169 | FALSE | TRUE | FALSE | FALSE |
| Myiagra rube | Aves | Extant | 0.013535703 | FALSE | TRUE | FALSE | FALSE |
| Neochmia te | Aves | Extant | 0.0095 | TRUE | FALSE | FALSE | FALSE |
| Neophema c | Aves | Extant | 0.0468 | TRUE | FALSE | FALSE | FALSE |
| Neophema e | Aves | Extant | 0.0471 | TRUE | FALSE | FALSE | FALSE |
| Ninox conniv | Aves | Extant | 0.65 | FALSE | TRUE | TRUE | FALSE |
| Ninox novaes | Aves | Extant | 0.177 | FALSE | TRUE | TRUE | FALSE |
| Ninox strenu | Aves | Extant | 1.35 | FALSE | FALSE | TRUE | FALSE |
| Northiella ha | Aves | Extant | 0.081820833 | TRUE | FALSE | FALSE | FALSE |
| Nycticorax ca | Aves | Extant | 0.725 | FALSE | TRUE | TRUE | TRUE |
| Nymphicus h | Aves | Extant | 0.092427796 | TRUE | FALSE | FALSE | FALSE |
| Ocyphaps lop | Aves | Extant | 0.192 | TRUE | FALSE | FALSE | FALSE |
| Oreoica gutt | Aves | Extant | 0.063418561 | TRUE | TRUE | FALSE | FALSE |
| Oriolus sagit | Aves | Extant | 0.097172222 | TRUE | TRUE | FALSE | FALSE |
| Oxyura austr | Aves | Extant | 0.801 | TRUE | FALSE | FALSE | TRUE |
| Pachycephala | Aves | Extant | 0.0308 | TRUE | TRUE | FALSE | FALSE |
| Pachycephala | Aves | Extant | 0.041006534 | TRUE | TRUE | FALSE | FALSE |
| Pachycephala | Aves | Extant | 0.0385 | TRUE | TRUE | FALSE | FALSE |
| Pachycephala | Aves | Extant | 0.0235 | TRUE | TRUE | FALSE | FALSE |
| Pachycephala | Aves | Extant | 0.036722222 | FALSE | TRUE | FALSE | FALSE |
| Pardalotus p | Aves | Extant | 0.0085 | TRUE | TRUE | FALSE | FALSE |

| | | | | | | | |
|---|---|---|---|---|---|---|---|
| Pardalotus st | Aves | Extant | 0.011 | FALSE | TRUE | FALSE | FALSE |
| Pedionomus | Aves | Extant | 0.0632 | TRUE | TRUE | FALSE | FALSE |
| Pelecanus co | Aves | Extant | 5.4 | FALSE | FALSE | TRUE | TRUE |
| Petrochelido | Aves | Extant | 0.0108 | FALSE | TRUE | FALSE | FALSE |
| Petrochelido | Aves | Extant | 0.0166 | FALSE | TRUE | FALSE | FALSE |
| Petroica goo | Aves | Extant | 0.0087 | FALSE | TRUE | FALSE | FALSE |
| Petroica pho | Aves | Extant | 0.0132 | FALSE | TRUE | FALSE | FALSE |
| Petroica rodi | Aves | Extant | 0.009698676 | FALSE | TRUE | FALSE | FALSE |
| Petroica rose | Aves | Extant | 0.008625 | FALSE | TRUE | FALSE | FALSE |
| Pezoporus w | Aves | Extant | 0.0826625 | TRUE | FALSE | FALSE | FALSE |
| Phalacrocora | Aves | Extant | 2.54 | FALSE | FALSE | FALSE | TRUE |
| Phalacrocora | Aves | Extant | 0.86 | FALSE | FALSE | FALSE | TRUE |
| Phalacrocora | Aves | Extant | 1.74 | FALSE | FALSE | FALSE | TRUE |
| Phaps chalco | Aves | Extant | 0.331 | TRUE | TRUE | FALSE | FALSE |
| Phaps elegar | Aves | Extant | 0.211 | TRUE | FALSE | FALSE | FALSE |
| Phylidonyris | Aves | Extant | 0.0209 | TRUE | TRUE | FALSE | FALSE |
| Platalea flav | Aves | Extant | 1.87 | FALSE | TRUE | FALSE | TRUE |
| Platalea regi | Aves | Extant | 1.73 | FALSE | TRUE | FALSE | TRUE |
| Platycercus a | Aves | Extant | 0.103 | TRUE | TRUE | FALSE | FALSE |
| Platycercus e | Aves | Extant | 0.126 | TRUE | TRUE | FALSE | FALSE |
| Platycercus e | Aves | Extant | 0.104 | TRUE | FALSE | FALSE | FALSE |
| Plegadis falc | Aves | Extant | 0.604 | TRUE | FALSE | FALSE | TRUE |
| Pluvialis fulv | Aves | Extant | 0.138 | FALSE | TRUE | FALSE | TRUE |
| Podargus str | Aves | Extant | 0.277 | FALSE | TRUE | TRUE | FALSE |
| Podiceps cris | Aves | Extant | 1.03 | FALSE | FALSE | FALSE | TRUE |
| Poliocephalu | Aves | Extant | 0.241 | FALSE | FALSE | FALSE | TRUE |
| Polytelis antl | Aves | Extant | 0.175 | TRUE | FALSE | FALSE | FALSE |
| Pomatostom | Aves | Extant | 0.0415 | TRUE | TRUE | TRUE | FALSE |
| Pomatostom | Aves | Extant | 0.0698 | TRUE | TRUE | FALSE | FALSE |
| Porphyrio po | Aves | Extant | 0.793 | TRUE | FALSE | FALSE | FALSE |

| | | | | | | | |
|---|---|---|---|---|---|---|---|
| Porzana flum | Aves | Extant | 0.064775 | TRUE | TRUE | FALSE | TRUE |
| Porzana pusi | Aves | Extant | 0.0339 | TRUE | FALSE | FALSE | TRUE |
| Porzana tabu | Aves | Extant | 0.0435 | TRUE | TRUE | FALSE | TRUE |
| Psephotus ha | Aves | Extant | 0.063 | TRUE | FALSE | FALSE | FALSE |
| Psophodes n | Aves | Extant | 0.0460125 | FALSE | TRUE | FALSE | FALSE |
| Purnella albi | Aves | Extant | 0.017233333 | TRUE | TRUE | FALSE | FALSE |
| Recurvirostra | Aves | Extant | 0.245 | FALSE | TRUE | FALSE | TRUE |
| Rhipidura leu | Aves | Extant | 0.0207 | FALSE | TRUE | FALSE | FALSE |
| Rhipidura ruf | Aves | Extant | 0.009669583 | FALSE | TRUE | FALSE | FALSE |
| Rostratula au | Aves | Extant | 0.127 | TRUE | FALSE | FALSE | TRUE |
| Sericornis fro | Aves | Extant | 0.0131 | TRUE | TRUE | FALSE | FALSE |
| Smicrornis b | Aves | Extant | 0.005725 | FALSE | TRUE | FALSE | FALSE |
| Stagonopleu | Aves | Extant | 0.0138 | TRUE | FALSE | FALSE | FALSE |
| Stagonopleu | Aves | Extant | 0.0178 | TRUE | FALSE | FALSE | FALSE |
| Stictonetta n | Aves | Extant | 0.912 | TRUE | FALSE | FALSE | TRUE |
| Stiltia isabel | Aves | Extant | 0.063655844 | TRUE | TRUE | FALSE | FALSE |
| Stipiturus ma | Aves | Extant | 0.0072 | FALSE | TRUE | FALSE | FALSE |
| Stipiturus ma | Aves | Extant | 0.0055 | TRUE | TRUE | FALSE | FALSE |
| Strepera gra | Aves | Extant | 0.308 | TRUE | TRUE | TRUE | FALSE |
| Strepera vers | Aves | Extant | 0.388 | TRUE | TRUE | TRUE | FALSE |
| Struthidea ci | Aves | Extant | 0.136 | TRUE | TRUE | TRUE | FALSE |
| Sugomel nig | Aves | Extant | 0.009256316 | TRUE | TRUE | FALSE | FALSE |
| Tachybaptus | Aves | Extant | 0.184 | FALSE | FALSE | FALSE | TRUE |
| Tadorna radj | Aves | Extant | 0.95 | TRUE | TRUE | FALSE | TRUE |
| Tadorna tado | Aves | Extant | 1.43 | TRUE | TRUE | FALSE | TRUE |
| Taeniopygia | Aves | Extant | 0.011128747 | TRUE | FALSE | FALSE | FALSE |
| Thalasseus b | Aves | Extant | 0.331 | FALSE | FALSE | FALSE | TRUE |
| Threskiornis | Aves | Extant | 1.35 | FALSE | TRUE | TRUE | FALSE |
| Todiramphus | Aves | Extant | 0.051728125 | FALSE | TRUE | TRUE | FALSE |
| Todiramphus | Aves | Extant | 0.053 | FALSE | TRUE | TRUE | TRUE |

| | | | | | | | |
|---|---|---|---|---|---|---|---|
| Tribonyx ven | Aves | Extant | 0.387 | TRUE | TRUE | FALSE | TRUE |
| Trichoglossu | Aves | Extant | 0.129 | TRUE | FALSE | FALSE | FALSE |
| Tringa glare | Aves | Extant | 0.05735 | FALSE | TRUE | FALSE | TRUE |
| Tringa nebul | Aves | Extant | 0.176 | FALSE | TRUE | FALSE | TRUE |
| Tringa stagn | Aves | Extant | 0.0702 | FALSE | TRUE | FALSE | TRUE |
| Turnix pyrrh | Aves | Extant | 0.04537601 | TRUE | TRUE | FALSE | FALSE |
| Turnix velox | Aves | Extant | 0.045 | TRUE | TRUE | FALSE | FALSE |
| Tyto alba | Aves | Extant | 0.392 | FALSE | FALSE | TRUE | FALSE |
| Tyto novaeh | Aves | Extant | 0.604 | FALSE | FALSE | TRUE | FALSE |
| Vanellus mil | Aves | Extant | 0.316 | FALSE | TRUE | FALSE | TRUE |
| Vanellus tric | Aves | Extant | 0.186 | TRUE | TRUE | FALSE | FALSE |
| Zoothera lun | Aves | Extant | 0.111 | FALSE | TRUE | FALSE | FALSE |
| Zosterops la | Aves | Extant | 0.011696519 | TRUE | TRUE | FALSE | FALSE |
| Ningaui yvon | Mammalia | Extant | 0.0082 | TRUE | TRUE | FALSE | FALSE |
| Bettongia pe | Mammalia | Extant | 1.26 | TRUE | TRUE | FALSE | FALSE |
| Lagorchestes | Mammalia | Extant | 3 | TRUE | FALSE | FALSE | FALSE |
| Leipoa gallin | Aves | Extinct | 5.5 | TRUE | TRUE | FALSE | FALSE |
| Megalibgwil | Mammalia | Extinct | 11 | FALSE | TRUE | FALSE | FALSE |
| Metasthenur | Mammalia | Extinct | 55 | TRUE | FALSE | FALSE | FALSE |
| Procoptodon | Mammalia | Extinct | 50 | TRUE | FALSE | FALSE | FALSE |
| Procoptodon | Mammalia | Extinct | 54 | TRUE | FALSE | FALSE | FALSE |
| Procoptodon | Mammalia | Extinct | 220 | TRUE | FALSE | FALSE | FALSE |
| Propleopus c | Mammalia | Extinct | 70 | TRUE | TRUE | TRUE | FALSE |
| Protemnodo | Mammalia | Extinct | 110 | TRUE | FALSE | FALSE | FALSE |
| Protemnodo | Mammalia | Extinct | 170 | TRUE | FALSE | FALSE | FALSE |
| Sarcophilus l | Mammalia | Extinct | 15 | FALSE | FALSE | TRUE | FALSE |
| Simosthenur | Mammalia | Extinct | 55 | TRUE | FALSE | FALSE | FALSE |
| Simosthenur | Mammalia | Extinct | 78 | TRUE | FALSE | FALSE | FALSE |
| Simosthenur | Mammalia | Extinct | 118 | TRUE | FALSE | FALSE | FALSE |
| Sthenurus ar | Mammalia | Extinct | 72 | TRUE | FALSE | FALSE | FALSE |

| | | | | | | | |
|---|---|---|---|---|---|---|---|
| Thylacoleo ca | Mammalia | Extinct | 115 | FALSE | FALSE | TRUE | FALSE |
| Wonambi na | Reptilia | Extinct | 30 | FALSE | FALSE | TRUE | FALSE |
| Zygomaturus | Mammalia | Extinct | 400 | TRUE | FALSE | FALSE | FALSE |
| Diprotodon o | Mammalia | Extinct | 2800 | TRUE | FALSE | FALSE | FALSE |
| Macropus fu | Mammalia | Extinct/Extant | 50 | TRUE | FALSE | FALSE | FALSE |

Table S2 Source of body mass and/or diet for extinct species

| SciName | Status | mean.mass.kg | eat.plant | eat.invert | eat.vert | eat.FishInv |
|---|---|---|---|---|---|---|
| **Leipoa gallinacea/Latigallina naracoortensis** | Extinct | 5.5 | TRUE | TRUE | FALSE | FALSE |

Boles, Walter E. "Systematics of the Fossil Australian Giant Megapodes Progura (Aves: Megapodiidae)" 7 (2008). Oryctos, 7, pp.195-215.
Van Tets, Gerard Frederick. "A revision of the fossil Megapodiidae (Aves), including a description of a new species of Progura De Vis." (1974).
Renema, Willem, ed. Biogeography, Time, and Place: Distributions, Barriers, and Islands. Vol. 29. Topics In Geobiology.
Dordrecht: Springer Netherlands, 2007. https://doi.org/10.1007/978-1-4020-6374-9.
Shute, Elen, Gavin J. Prideaux, and Trevor H. Worthy. "Taxonomic Review of the Late Cenozoic Megapodes (Galliformes: Megapodiidae) of Australia."
Royal Society Open Science 4, no. 6 (June 2017): 170233. https://doi.org/10.1098/rsos.170233.

| SciName | Status | mean.mass.kg | eat.plant | eat.invert | eat.vert | eat.FishInv |
|---|---|---|---|---|---|---|
| **Megalibgwilia ramsayi** | Extinct | 11 | FALSE | TRUE | FALSE | FALSE |

Ashwell, Ken W.S., Craig D. Hardman, and Anne M. Musser. "Brain and Behaviour of Living and Extinct Echidnas." Zoology 117, no. 5 (October 2014): 349–61. https://doi.org/10.1016/j.zool.2014.05.002.
Johnson, Chris. Australia's Mammal Extinctions: A 50,000-Year History. Cambridge University Press, 2006.

| SciName | Status | mean.mass.kg | eat.plant | eat.invert | eat.vert | eat.FishInv |
|---|---|---|---|---|---|---|
| **Metasthenurus newtonae** | Extinct | 55 | TRUE | FALSE | FALSE | FALSE |

Johnson, Chris. Australia's Mammal Extinctions: A 50,000-Year History. Cambridge University Press, 2006.

| SciName | Status | mean.mass.kg | eat.plant | eat.invert | eat.vert | eat.FishInv |
|---|---|---|---|---|---|---|
| **Palorchestes azael** | Extinct | 500 | TRUE | FALSE | FALSE | FALSE |

Johnson, Chris. Australia's Mammal Extinctions: A 50,000-Year History. Cambridge University Press, 2006.

| SciName | Status | mean.mass.kg | eat.plant | eat.invert | eat.vert | eat.FishInv |
|---|---|---|---|---|---|---|
| **Procoptodon browneorum** | Extinct | 50 | TRUE | FALSE | FALSE | FALSE |

Johnson, Chris. Australia's Mammal Extinctions: A 50,000-Year History. Cambridge University Press, 2006.

| SciName | Status | mean.mass.kg | eat.plant | eat.invert | eat.vert | eat.FishInv |
|---|---|---|---|---|---|---|
| **Procoptodon gilli** | Extinct | 54 | TRUE | FALSE | FALSE | FALSE |

Johnson, Chris. Australia's Mammal Extinctions: A 50,000-Year History. Cambridge University Press, 2006.

| SciName | Status | mean.mass.kg | eat.plant | eat.invert | eat.vert | eat.FishInv |
|---|---|---|---|---|---|---|

| | | | | | | |
|---|---|---|---|---|---|---|
| **Procoptodon goliah** | Extinct | 220 | TRUE | FALSE | FALSE | FALSE |

Johnson, Chris. Australia's Mammal Extinctions: A 50,000-Year History. Cambridge University Press, 2006.

Helgen, Kristofer M., Rod T. Wells, Benjamin P. Kear, Wayne R. Gerdtz, and Timothy F. Flannery. "Ecological and Evolutionary Significance of Sizes of Giant Extinct Kangaroos." Australian Journal of Zoology 54, no. 4 (2006): 293. https://doi.org/10.1071/ZO

Janis, Christine M., Karalyn Buttrill, and Borja Figueirido. "Locomotion in Extinct Giant Kangaroos: Were Sthenurines Hop-Less Monsters?" Edited by Brian Lee Beatty. PLoS ONE 9, no. 10 (October 15, 2014): e109888.

| **SciName** | **Status** | **mean.mass.kg** | **eat.plant** | **eat.invert** | **eat.vert** | **eat.FishInv** |
|---|---|---|---|---|---|---|
| **Propleopus oscillans** | Extinct | 70 | TRUE | TRUE | TRUE | FALSE |

Wroe, Stephen, Christine Argot, and Christopher Dickman. "On the Rarity of Big Fierce Carnivores and Primacy of Isolation and Area: Tracking Large Mammalian Carnivore Diversity on Two Isolated Continents." Proceedings of the Royal Society of London Series B: Biological Sciences. 2004 Jun 7;271(1544):1203-11.

Johnson, Chris. Australia's Mammal Extinctions: A 50,000-Year History. Cambridge University Press, 2006.

Ride, William David Lindsay, et al. "Towards a biology of Propleopus oscillans (Marsupialia: Propleopinae, Hypsiprymnodontidae)." PROCEEDINGS-LINNEAN SOCIETY OF NEW SOUTH WALES. Vol. 117. LINNEAN SOCIETY OF NEW SOUTH WALES, 1997.

Johnson, Chris N. "The rise and fall of large marsupial carnivores." A. Glen, D., and Dickman, eds. Carnivores of Australia, past, present and future. CSIRO, Collingwood, Australia (2014): 13-26.

| **SciName** | **Status** | **mean.mass.kg** | **eat.plant** | **eat.invert** | **eat.vert** | **eat.FishInv** |
|---|---|---|---|---|---|---|
| **Protemnodon brehus** | Extinct | 110 | TRUE | FALSE | FALSE | FALSE |

Johnson, Chris. Australia's Mammal Extinctions: A 50,000-Year History. Cambridge University Press, 2006.

| **SciName** | **Status** | **mean.mass.kg** | **eat.plant** | **eat.invert** | **eat.vert** | **eat.FishInv** |
|---|---|---|---|---|---|---|
| **Protemnodon roechus** | Extinct | 170 | TRUE | FALSE | FALSE | FALSE |

Johnson, Chris. Australia's Mammal Extinctions: A 50,000-Year History. Cambridge University Press, 2006.

| **SciName** | **Status** | **mean.mass.kg** | **eat.plant** | **eat.invert** | **eat.vert** | **eat.FishInv** |
|---|---|---|---|---|---|---|
| **Sarcophilus laniarius** | Extinct | 17.8 (Wroe has | FALSE | FALSE | TRUE | FALSE |

Wroe, S. "A Review of Terrestrial Mammalian and Reptilian Carnivore Ecology in Australian Fossil Faunas, and Factors

Influencing Their Diversity: The Myth of Reptilian Domination and Its Broader Ramifications." Australian Journal of Zoology 50 (1): 1-24

Rose, Robert K., David A. Pemberton, Nick J. Mooney, and Menna E. Jones. "Sarcophilus Harrisii (Dasyuromorphia: Dasyuridae)." Mammalian Species 49, no. 942 (May 1, 2017): 1–17. https://doi.org/10.1093/mspecies/sex001.

| **SciName** | **Status** | **mean.mass.kg** | **eat.plant** | **eat.invert** | **eat.vert** | **eat.FishInv** |
|---|---|---|---|---|---|---|
| **Simosthenurus baileyi** | Extinct | 55 | TRUE | FALSE | FALSE | FALSE |

Johnson, Chris. Australia's Mammal Extinctions: A 50,000-Year History. Cambridge University Press, 2006.

| **SciName** | **Status** | **mean.mass.kg** | **eat.plant** | **eat.invert** | **eat.vert** | **eat.FishInv** |
|---|---|---|---|---|---|---|
| **Simosthenurus maddocki** | Extinct | 78 | TRUE | FALSE | FALSE | FALSE |

Johnson, Chris. Australia's Mammal Extinctions: A 50,000-Year History. Cambridge University Press, 2006.

| **SciName** | **Status** | **mean.mass.kg** | **eat.plant** | **eat.invert** | **eat.vert** | **eat.FishInv** |
|---|---|---|---|---|---|---|
| **Simosthenurus occidentalis** | Extinct | 118 | TRUE | FALSE | FALSE | FALSE |

Johnson, Chris. Australia's Mammal Extinctions: A 50,000-Year History. Cambridge University Press, 2006.

| **SciName** | **Status** | **mean.mass.kg** | **eat.plant** | **eat.invert** | **eat.vert** | **eat.FishInv** |
|---|---|---|---|---|---|---|
| **Simosthenurus pales** | Extinct | 150 | TRUE | FALSE | FALSE | FALSE |

Johnson, Chris. Australia's Mammal Extinctions: A 50,000-Year History. Cambridge University Press, 2006.

| **SciName** | **Status** | **mean.mass.kg** | **eat.plant** | **eat.invert** | **eat.vert** | **eat.FishInv** |
|---|---|---|---|---|---|---|
| **Sthenurus andersoni** | Extinct | 72 | TRUE | FALSE | FALSE | FALSE |

Johnson, Chris. Australia's Mammal Extinctions: A 50,000-Year History. Cambridge University Press, 2006.

| **SciName** | **Status** | **mean.mass.kg** | **eat.plant** | **eat.invert** | **eat.vert** | **eat.FishInv** |
|---|---|---|---|---|---|---|
| **Thylacoleo carnifex** | Extinct | 115 | FALSE | FALSE | TRUE | FALSE |

Johnson, Chris. Australia's Mammal Extinctions: A 50,000-Year History. Cambridge University Press, 2006.

| **SciName** | **Status** | **mean.mass.kg** | **eat.plant** | **eat.invert** | **eat.vert** | **eat.FishInv** |
|---|---|---|---|---|---|---|
| **Wonambi naracoortensis** | Extinct | 30 (big discrepe | FALSE | FALSE | TRUE | FALSE |

Wroe, S. "A Review of Terrestrial Mammalian and Reptilian Carnivore Ecology in Australian Fossil Faunas, and Factors

Influencing Their Diversity: The Myth of Reptilian Domination and Its Broader Ramifications.” Australian Journal of Zoology 50 (1): 1-24.
Scanlon, John D. "Giant terrestrial reptilian carnivores of Cenozoic Australia." Carnivores of Australia: Past, Present and Future (2014): 27.

| SciName | Status | mean.mass.kg | eat.plant | eat.invert | eat.vert | eat.FishInv |
|---|---|---|---|---|---|---|
| **Zygomaturus trilobus** | Extinct | 500 | TRUE | FALSE | FALSE | FALSE |

Johnson, Chris. Australia’s Mammal Extinctions: A 50,000-Year History. Cambridge University Press, 2006.
Gillespie, Richard, Aaron B. Camens, Trevor H. Worthy, Nicolas J. Rawlence, Craig Reid, Fiona Bertuch, Vladimir Levchenko, and Alan Cooper. “Man and Megafauna in Tasmania: Closing the Gap.” Quaternary Science Reviews 37 (March 2012): 38–47. https://doi.or
Sharp, Alana C. “A Quantitative Comparative Analysis of the Size of the Frontoparietal Sinuses and Brain in Vombatiform Marsupials.” Memoirs of Museum Victoria 74 (2016): 331–42. https://doi.org/10.24199/j.mmv.2016.74.23.
Murray, Peter. "The Pleistocene megafauna of Australia." *Vertebrate palaeontology of Australasia* (1991): 1070-1164.

| SciName | Status | mean.mass.kg | eat.plant | eat.invert | eat.vert | eat.FishInv |
|---|---|---|---|---|---|---|
| **Diprotodon optatum** | Extinct | 2700 | TRUE | FALSE | FALSE | FALSE |

Johnson, Chris. Australia’s Mammal Extinctions: A 50,000-Year History. Cambridge University Press, 2006.
Wroe, Stephen, Mathew Crowther, Joe Dortch, and John Chong. “The Size of the Largest Marsupial and Why It Matters.” Proceedings of the Royal Society of London. Series B: Biological Sciences 271, no. suppl_3 (February 7,2004). https://doi.org/10.1098/rsbl

| SciName | Status | mean.mass.kg | eat.plant | eat.invert | eat.vert | eat.FishInv |
|---|---|---|---|---|---|---|
| **Macropus fuliginosus/giganteus/titan** | Extinct/Extant | 50 | TRUE | FALSE | FALSE | FALSE |

Janis, Christine M., Karalyn Buttrill, and Borja Figueirido. “Locomotion in Extinct Giant Kangaroos: Were Sthenurines Hop-Less Monsters?” Edited by Brian Lee Beatty. PLoS ONE 9, no. 10 (October 15, 2014): e109888. https://doi.org/10.1371/journal.pone.0109
Helgen, Kristofer M., Rod T. Wells, Benjamin P. Kear, Wayne R. Gerdtz, and Timothy F. Flannery. “Ecological and Evolutionary Significance of Sizes of Giant Extinct Kangaroos.” Australian Journal of Zoology 54, no. 4 (2006): 293. https://doi.org/10.1071/ZO

**General references (details for multiple species)**

Smith, Felisa A., Rosemary E. Elliott Smith, S. Kathleen Lyons, and Jonathan L. Payne. “Body Size Downgrading of Mammals over the Late Quaternary.” Science 360, no. 6386 (April 20, 2018): 310–13. https://doi.org/10.1126/science.aao5987.
Flannery, T. F. “Pleistocene Faunal Loss: Implications of the Aftershock for Australia’s Past and Future.” Archaeology in Oceania 25, no. 2 (July 1990): 45–55. https://doi.org/10.1002/j.1834-4453.1990.tb00232.x.

Johnson, Chris. Australia's Mammal Extinctions: A 50,000-Year History. Cambridge University Press, 2006.
Camens, Aaron B. Systematic and palaeobiological implications of postcranial morphology in the Diprotodontidae (Marsupialia). Doctoral dissertation (2010).

Table S3 fist page of interaction data from GloBI

| resource | consumer | body_mass_resource | body_mass_consumer | class_resource | class_consumer | endo/ecto |
|---|---|---|---|---|---|---|
| Gallus gallus | Bubo virginianus | 751.72 | 1575.7 | aves | aves | endotherm |
| Mus musculus | Bubo virginianus | 17.775 | 1575.7 | mammals | aves | endotherm |
| Megascops asio | Bubo virginianus | 179.99 | 1575.7 | aves | aves | endotherm |
| Rattus norvegicus | Bubo virginianus | 310.665 | 1575.7 | mammals | aves | endotherm |
| Cyanocitta cristata | Bubo virginianus | 88 | 1575.7 | aves | aves | endotherm |
| Corvus brachyrhynchos | Bubo virginianus | 448.76 | 1575.7 | aves | aves | endotherm |
| Bonasa umbellus | Bubo virginianus | 530.91 | 1575.7 | aves | aves | endotherm |
| Columba livia | Bubo virginianus | 354.2 | 1575.7 | aves | aves | endotherm |
| Sciurus niger | Bubo virginianus | 761.9 | 1575.7 | mammals | aves | endotherm |
| Plectrophenax nivalis | Bubo virginianus | 42.2 | 1575.7 | aves | aves | endotherm |
| Aythya marila | Bubo virginianus | 1005.37 | 1575.7 | aves | aves | endotherm |
| Gallinula chloropus | Bubo virginianus | 339.63 | 1575.7 | aves | aves | endotherm |
| Turdus migratorius | Bubo virginianus | 79.98562144 | 1575.7 | aves | aves | endotherm |
| Agelaius phoeniceus | Bubo virginianus | 50.78 | 1575.7 | aves | aves | endotherm |
| Melospiza melodia | Bubo virginianus | 21.91 | 1575.7 | aves | aves | endotherm |
| Euphagus carolinus | Bubo virginianus | 59.57 | 1575.7 | aves | aves | endotherm |
| Ondatra zibethicus | Bubo virginianus | 1028.875 | 1575.7 | mammals | aves | endotherm |
| Podilymbus podiceps | Bubo virginianus | 411.93 | 1575.7 | aves | aves | endotherm |
| Anas platyrhynchos | Bubo virginianus | 843.42 | 1575.7 | aves | aves | endotherm |
| Tringa flavipes | Bubo virginianus | 77.5 | 1575.7 | aves | aves | endotherm |
| Asio otus | Bubo virginianus | 296.57 | 1575.7 | aves | aves | endotherm |
| Rallus elegans | Bubo virginianus | 310.6604002 | 1575.7 | aves | aves | endotherm |
| Sciurus carolinensis | Strix varia | 526.25 | 711.49 | mammals | aves | endotherm |
| Megascops asio | Strix varia | 179.99 | 711.49 | aves | aves | endotherm |
| Bonasa umbellus | Strix varia | 530.91 | 711.49 | aves | aves | endotherm |
| Cyanocitta cristata | Strix varia | 88 | 711.49 | aves | aves | endotherm |
| Turdus migratorius | Strix varia | 79.98562144 | 711.49 | aves | aves | endotherm |
| Mus musculus | Strix varia | 17.775 | 711.49 | mammals | aves | endotherm |
| Sciurus niger | Strix varia | 761.9 | 711.49 | mammals | aves | endotherm |

**Supplementary Table S4 Quantile regression models tested**

| **Quantile regression** | **BIC values for 0.95 quantile** | **BIC values for 0.05 quantile** |
|---|---|---|
| prey mass~1 | 16974.37 | 12401.17 |
| prey mass~predator class | 15546.87 | 12315.05 |
| prey mass ~predator mass | 13941.17 | 11566.74 |
| prey mass~predator mass+predator class | 13942.66 | **11272.55** |
| prey mass ~predatormass*predator class | **13868.49** | 11274.53 |

**Table S5 True skill statistics for the new trophic niche space model (that includes different predator/prey body size relationships depending on predator class) *versus* the simple body size trophic niche space model. In addition to the true skill statistic, the table includes the number of true positives (a), true negatives (d), false positives (b), and false negatives (c). Whilst the false positives are quite high, some of these false positives are likely true positives that have not been observed (due to incomplete data).**

| model | data | TSS | a | b | c | d |
|---|---|---|---|---|---|---|
| new model | GloBI | 0.384 | 864 | 494 | 99 | 469 |
| simple body size model | GloBI | 0.372 | 864 | 506 | 99 | 457 |
| New model | Serengeti | 0.595 | 184 | 869 | 17 | 1846 |
| simple body size model | Serengeti | 0.579 | 183 | 900 | 18 | 1815 |

**Table S6a Diet breadths from the literature**

| species | breadth | eat plants | eat inverts | eat verts | eat fish | source |
|---|---|---|---|---|---|---|
| *Hoplocephalus bungaroides* | 6 | FALSE | FALSE | TRUE | FALSE | [1] |
| *Python reticulatus* | 7 | FALSE | FALSE | TRUE | FALSE | [2] |
| *Panthera pardus* | 8 | FALSE | FALSE | TRUE | FALSE | [3] |
| *Pseudonaja affinis* | 11 | FALSE | FALSE | TRUE | FALSE | [4] |
| *Panthera tigris* | 12 | FALSE | FALSE | TRUE | FALSE | [5] |
| *Panthera pardus* | 14 | FALSE | FALSE | TRUE | FALSE | [5] |
| *Varanus varius* | 16 | FALSE | TRUE | TRUE | FALSE | [6] |
| *Ninox rufa* | 17 | FALSE | TRUE | TRUE | FALSE | [7] |
| *Canis lupus* | 20 | FALSE | FALSE | TRUE | FALSE | [8] |
| *Ninox strenua* | 26 | FALSE | FALSE | TRUE | FALSE | [9] |
| *Canis lupus* | 28 | FALSE | FALSE | TRUE | FALSE | [10] |
| *Dasyurus maculatus* | 29 | FALSE | TRUE | TRUE | FALSE | [11] |
| *Vulpes vulpes* | 31 | TRUE | TRUE | TRUE | FALSE | [10] |
| *Hieraaetus morphnoides* | 36 | FALSE | TRUE | TRUE | FALSE | [12] |
| *Hieraaetus morphnoides* | 46 | FALSE | TRUE | TRUE | FALSE | [12] |
| *Aquila audax* | 70 | FALSE | FALSE | TRUE | FALSE | [12] |
| *Tropidonophis mairii* | 10 | FALSE | FALSE | TRUE | FALSE | [13] |
| *Haliastur sphenurus* | 26 | FALSE | TRUE | TRUE | TRUE | [14] |
| *Falco subniger* | 10 | FALSE | TRUE | TRUE | FALSE | [14] |
| *Ninox connivens* | 11 | FALSE | TRUE | TRUE | FALSE | [14] |
| *Redunca redunca* | 3 | TRUE | FALSE | FALSE | FALSE | [15] |
| *Ourebia ourebi* | 4 | TRUE | FALSE | FALSE | FALSE | “ |
| *Tragelaphus scriptus* | 4 | TRUE | FALSE | FALSE | FALSE | “ |

| | | | | | | |
|---|---|---|---|---|---|---|
| *Pedetes capensis* | 8 | TRUE | FALSE | FALSE | FALSE | “ |
| *Hippopotamus amphibius* | 9 | TRUE | FALSE | FALSE | FALSE | “ |
| *Kobus ellipsiprymnus* | 12 | TRUE | FALSE | FALSE | FALSE | “ |
| *Damaliscus korrigum* | 14 | TRUE | FALSE | FALSE | FALSE | “ |
| *Taurotragus oryx* | 15 | TRUE | FALSE | FALSE | FALSE | “ |
| *Syncerus caffer* | 18 | TRUE | FALSE | FALSE | FALSE | “ |
| *Equus quagga* | 19 | TRUE | FALSE | FALSE | FALSE | “ |
| *Giraffa camelopardalis* | 20 | TRUE | FALSE | FALSE | FALSE | “ |
| *Alcelaphus buselaphus* | 21 | TRUE | FALSE | FALSE | FALSE | “ |
| *Aepyceros melampus* | 23 | TRUE | FALSE | FALSE | FALSE | “ |
| *Eudorcas thomsonii* | 25 | TRUE | FALSE | FALSE | FALSE | “ |
| *Connochaetes taurinus* | 26 | TRUE | FALSE | FALSE | FALSE | “ |
| *Nanger granti* | 27 | TRUE | FALSE | FALSE | FALSE | “ |
| *Madoqua kirkii* | 30 | TRUE | FALSE | FALSE | FALSE | “ |
| *Loxodonta africana* | 46 | TRUE | FALSE | FALSE | FALSE | “ |
| *Heterohyrax brucei* | 61 | TRUE | FALSE | FALSE | FALSE | “ |
| *Procavia capensis* | 73 | TRUE | FALSE | FALSE | FALSE | [15] |
| *Hirundo rustica* | 76 | TRUE | TRUE | FALSE | FALSE | [16] |
| *Tachyglossus aculeatus* | 76 | FALSE | TRUE | FALSE | FALSE | [17] |
| *Apus apus* | 78 | FALSE | TRUE | FALSE | FALSE | [16] |
| *Delichon urbicum* | 113 | FALSE | TRUE | FALSE | FALSE | [16] |
| *Pteronotus personatus* | 114 | FALSE | TRUE | FALSE | FALSE | [18] |
| *Pteronotus davyi* | 169 | FALSE | TRUE | FALSE | FALSE | [18] |
| *Pteronotus parnelli* | 198 | FALSE | TRUE | FALSE | FALSE | [18] |
| *Perognathus formosus* | 27 | TRUE | TRUE | FALSE | FALSE | [19] |
| *Morus bassanus* | 5 | FALSE | FALSE | FALSE | TRUE | [20] |
| *Phalacrocorax auritus* | 16 | FALSE | FALSE | FALSE | TRUE | [21] |

| | | | | | | |
|---|---|---|---|---|---|---|
| *Larus cachinnans* | 18 | FALSE | TRUE | TRUE | TRUE | [21] |
| *Ardea modesta* | 8 | FALSE | TRUE | TRUE | TRUE | [22] |
| *Ardea intermedia* | 9 | FALSE | TRUE | TRUE | TRUE | [22] |
| *Haliaeetus leucogaster* | 11 | FALSE | FALSE | TRUE | TRUE | [23] |
| *Phalacrocorax sulcirostris* | 13 | FALSE | TRUE | FALSE | TRUE | [24] |
| *Phalacrocorax varius* | 14 | FALSE | TRUE | FALSE | TRUE | [24] |
| *Haliaeetus leucogaster* | 33 | FALSE | FALSE | TRUE | TRUE | [14] |
| *Bitis arietans* | 32 | FALSE | FALSE | TRUE | FALSE | [25] |

17. Griffiths M, Greenslade PJM. 1990 The diet of the spiny-anteater Tachyglossus Aculeatus acanthion in tropical habitats in the Northern Territory. *Beagle: Records of the Museums and Art Galleries of the Northern Territory, The*

23. Debus SJS. 2008 Biology and Diet of the White-bellied Sea-eagle Haliaeetus leucogaster Breeding in Northern Inland New South Wales. *Australian Field Ornithology*

24. Miller B. 1979 Ecology of the Little Black Cormorant, Phalacrocorax sulcirostris, and Little Pied Cormorant, P. melanoleucos, in Inland New South Wales I. Food and Feeding Habits. *Wildl. Res.* **6**, 79–95. (doi:10.1071/wr9790079)

25. In press. Museum Specimens Bias Measures of Snake Diet: A Case Study Using the Ambush-Foraging Puff Adder (Bitis arietans) | Herpetologica. See https://meridian.allenpress.com/herpetologica/article-abstract/73/2/121/32918/Museum-Specimens-Bias-Measures-of-Snake-Diet-A (accessed on 5 July 2021).

**Table S6b fish diversity in the diets of Australian inland piscivores**

| species | breadth | fish diversity | eat plants | eat invertebrates | eat vertebrates | eat fish | source |
|---|---|---|---|---|---|---|---|
| Ardea modesta | 8 | 3 | FALSE | TRUE | TRUE | TRUE | [1] |
| Ardea intermedia | 9 | 1 | FALSE | TRUE | TRUE | TRUE | [1] |
| Haliaeetus leucogaster | 11 | 3 | FALSE | FALSE | TRUE | TRUE | [2] |
| Haliaeetus leucogaster | 23 | 4 | FALSE | FALSE | TRUE | TRUE | [3] |
| Phalacrocorax sulcirostris | 13 | 8 | FALSE | TRUE | FALSE | TRUE | [4] |
| Phalacrocorax varius | 14 | 8 | FALSE | TRUE | FALSE | TRUE | [4] |
| Chelodina Longicollis | ? | 2 | FALSE | TRUE | TRUE | TRUE | [5] |
| Hydromys chrysogaster | ? | 4 | FALSE | TRUE | TRUE | TRUE | [6] |
| Haliaeetus leucogaster | 33 | 6 | FALSE | FALSE | TRUE | TRUE | [7] |

1. Taylor IR, Schultz MC. 2008 Breeding Season Diets of Egrets in Southeast Australia: Implications for Future Population Viability. *cowa* **31**, 593–601. (doi:10.1675/1524-4695-31.4.593)

2. Debus SJS. 2008 Biology and Diet of the White-bellied Sea-eagle Haliaeetus leucogaster Breeding in Northern Inland New South Wales. *Australian Field Ornithology*

### Table S7 Network position metrics

**Trophic level**
Trophic level describes a species' position in its trophic network. Primary producers, such as plants, have a trophic level of 1, while consumers have a trophic level equal to that of the species they consume plus 1. Thus, if a species feeds from multiple trophic levels, its trophic level is calculated by taking the average trophic level of all the species it consumes and adding 1.
Reference:
Lindeman, R.L. The trophic-dynamic aspect of ecology. *Ecology* **23.4,** 399-417 (1942).

**Degree**
A node's degree is based on the number of links it has. Node degree is usually expressed in its normalized form (number of links divided by the number of links the node with the most links has).
Reference:
Freeman, L. C. Centrality in social networks conceptual clarification. *Social Networks* 1, 215 239 (1978).

**Eccentricity**
Eccentricity is the shortest path between the focal node and the node furthest from the focal node.
Reference:
Harary, F. Graph theory. Addison Wesley Publishing Company, First Edition (1969)

**Eigenvector centrality**
Eigenvector centrality measures how influential a node is based on the number of links it has, with these links weighted depending on the number of links the contributing nodes have i.e., links coming from nodes with many links have a higher weighting than links from nodes with few links.
Reference:
Bonacich, P. Power and Centrality: A Family of Measures. *American Journal of Sociology* 92, 1170–1182 (1987).

**PageRank**
PageRank score is a centrality similar to Eigenvector centrality. It is calculated based on the number of 'in' links a node (or page) receives, weighted depending on how many 'in' and 'out' links the contributing nodes have. Having more 'in' links adds weight to a node's 'out' links, but the weight of these 'out' links decrease as their number increases.
Reference:
Brin, S. & Page, L. The anatomy of a large-scale hypertextual Web search engine. *Computer Networks and ISDN Systems* **30**, 107–117 (1998).

**Betweenness centrality**
Betweenness centrality indicates how important a node is as a connector between other nodes. It is based on the number of shortest paths between nodes that travel through the focal node.
Reference:

Freeman, L. C. A Set of Measures of Centrality Based on Betweenness. *Sociometry* **40**, 35 (1977).

**Closeness centrality**

A node's closeness centrality is based on the path length from the focal node to all other nodes in the network. It is calculated by taking the inverse of the sum of shortest path lengths. However, it is often expressed in its normalized form, where the numerator is the number of nodes in the network minus 1, and the denominator is the sum of shortest paths.

**Coreness**

Coreness is a centrality measure that indicates which core a node belongs to. First, nodes with no links are removed. These nodes belong to core 0. Then, nodes with one link (or less) are removed (including nodes that started with more than one link, but ended up having one or no links after other nodes were removed). These nodes belong to core 1. This process is continued until a further step would result in no nodes remaining. The innermost core is known as the k-core.

**Table S8 Mixed-effects models of vulnerabilities calculated using the simulation method, including coefficient estimates, AICc and AICc weights, and conditional and marginal $r^2$**

| intercept | basal connections | diet breadth | trophic level | basal connections* diet breadth | basal connections* trophic level | diet breadth* trophic level | basal connections* diet breadth* trophic level | df | logLik | AICc | delta | weight | conditional $r^2$ | marginal $r^2$ |
|---|---|---|---|---|---|---|---|---|---|---|---|---|---|---|
| 0.460 | -0.495 | -0.111 | -0.075 | 0.078 | 0.082 | 0.081 | -0.058 | 10 | 2624 | -5229 | 0 | 1.000 | 0.99 | 0.55 |
| 0.417 | -0.433 | -0.058 | -0.029 | 0.024 | 0.017 | 0.026 | NA | 9 | 2608 | -5198 | 31 | 0.000 | 0.99 | 0.55 |
| 0.414 | -0.414 | -0.073 | -0.026 | 0.024 | NA | 0.039 | NA | 8 | 2607 | -5197 | 31 | 0.000 | 0.99 | 0.55 |
| 0.412 | -0.456 | -0.027 | -0.025 | 0.023 | 0.036 | NA | NA | 8 | 2605 | -5194 | 35 | 0.000 | 0.99 | 0.55 |
| 0.389 | -0.417 | -0.025 | NA | 0.016 | NA | NA | NA | 6 | 2590 | -5169 | 60 | 0.000 | 0.99 | 0.55 |
| 0.387 | -0.418 | -0.025 | 0.002 | 0.017 | NA | NA | NA | 7 | 2591 | -5167 | 62 | 0.000 | 0.99 | 0.55 |
| 0.400 | -0.432 | NA | -0.022 | NA | 0.021 | NA | NA | 6 | 2576 | -5140 | 89 | 0.000 | 0.99 | 0.55 |
| 0.399 | -0.428 | -0.002 | -0.020 | NA | 0.018 | NA | NA | 7 | 2577 | -5139 | 90 | 0.000 | 0.99 | 0.55 |
| 0.399 | -0.407 | -0.023 | -0.020 | NA | NA | 0.018 | NA | 7 | 2577 | -5139 | 90 | 0.000 | 0.99 | 0.55 |
| 0.400 | -0.418 | -0.014 | -0.021 | NA | 0.010 | 0.010 | NA | 8 | 2577 | -5138 | 91 | 0.000 | 0.99 | 0.55 |
| 0.383 | -0.412 | -0.005 | NA | NA | NA | NA | NA | 5 | 2572 | -5133 | 95 | 0.000 | 0.99 | 0.55 |
| 0.386 | -0.411 | -0.005 | -0.005 | NA | NA | NA | NA | 6 | 2573 | -5133 | 96 | 0.000 | 0.99 | 0.55 |
| 0.381 | -0.415 | NA | NA | NA | NA | NA | NA | 4 | 2569 | -5130 | 99 | 0.000 | 0.99 | 0.55 |
| 0.385 | -0.413 | NA | -0.005 | NA | NA | NA | NA | 5 | 2570 | -5130 | 99 | 0.000 | 0.99 | 0.55 |
| 0.275 | NA | -0.163 | -0.168 | NA | NA | 0.123 | NA | 6 | 1621 | -3229 | 1999 | 0.000 | 0.57 | 0.11 |
| 0.179 | NA | -0.041 | -0.071 | NA | NA | NA | NA | 5 | 1580 | -3149 | 2080 | 0.000 | 0.55 | 0.07 |
| 0.112 | NA | -0.043 | NA | NA | NA | NA | NA | 4 | 1542 | -3077 | 2152 | 0.000 | 0.55 | 0.05 |
| 0.155 | NA | NA | -0.075 | NA | NA | NA | NA | 4 | 1538 | -3067 | 2161 | 0.000 | 0.55 | 0.03 |
| 0.084 | NA | NA | NA | NA | NA | NA | NA | 3 | 1499 | -2992 | 2237 | 0.000 | 0.55 | 0.00 |

**Table S9 Mixed-effects models of vulnerabilities calculated using the Bayesian network method, including coefficient estimates, AICc and AICc weights, and conditional and marginal $r^2$**

| intercept | basal connections | diet breadth | trophic level | basal connections* diet breadth | basal connections* trophic level | diet breadth* trophic level | basal connections* diet breadth* trophic level | df | logLik | AICc | delta | weight | conditional $r^2$ | marginal $r^2$ |
|---|---|---|---|---|---|---|---|---|---|---|---|---|---|---|
| 0.170 | -0.155 | -0.065 | -0.043 | 0.047 | 0.049 | 0.047 | -0.035 | 10 | 4021 | -8021 | 0 | 1.000 | 0.94 | 0.69 |
| 0.145 | -0.118 | -0.033 | -0.016 | 0.014 | 0.010 | 0.014 | NA | 9 | 3996 | -7973 | 48 | 0.000 | 0.94 | 0.69 |
| 0.143 | -0.108 | -0.042 | -0.014 | 0.014 | NA | 0.022 | NA | 8 | 3994 | -7972 | 49 | 0.000 | 0.94 | 0.69 |
| 0.142 | -0.131 | -0.016 | -0.013 | 0.014 | 0.020 | NA | NA | 8 | 3992 | -7968 | 53 | 0.000 | 0.94 | 0.69 |
| 0.130 | -0.109 | -0.015 | NA | 0.010 | NA | NA | NA | 6 | 3972 | -7932 | 89 | 0.000 | 0.95 | 0.69 |
| 0.128 | -0.110 | -0.015 | 0.002 | 0.010 | NA | NA | NA | 7 | 3973 | -7931 | 90 | 0.000 | 0.95 | 0.69 |
| 0.135 | -0.117 | NA | -0.011 | NA | 0.011 | NA | NA | 6 | 3948 | -7883 | 138 | 0.000 | 0.95 | 0.68 |
| 0.134 | -0.115 | -0.001 | -0.010 | NA | 0.009 | NA | NA | 7 | 3948 | -7883 | 138 | 0.000 | 0.95 | 0.68 |
| 0.134 | -0.104 | -0.012 | -0.010 | NA | NA | 0.009 | NA | 7 | 3948 | -7882 | 139 | 0.000 | 0.95 | 0.68 |
| 0.135 | -0.111 | -0.006 | -0.011 | NA | 0.006 | 0.005 | NA | 8 | 3949 | -7881 | 140 | 0.000 | 0.95 | 0.68 |
| 0.126 | -0.107 | -0.002 | NA | NA | NA | NA | NA | 5 | 3943 | -7876 | 145 | 0.000 | 0.95 | 0.68 |
| 0.128 | -0.106 | -0.002 | -0.002 | NA | NA | NA | NA | 6 | 3944 | -7876 | 146 | 0.000 | 0.95 | 0.68 |
| 0.126 | -0.108 | NA | NA | NA | NA | NA | NA | 4 | 3940 | -7872 | 149 | 0.000 | 0.95 | 0.68 |
| 0.127 | -0.107 | NA | -0.002 | NA | NA | NA | NA | 5 | 3941 | -7872 | 149 | 0.000 | 0.95 | 0.68 |
| 0.103 | NA | -0.053 | -0.047 | NA | NA | 0.039 | NA | 6 | 3394 | -6776 | 1245 | 0.000 | 0.65 | 0.08 |
| 0.073 | NA | -0.014 | -0.017 | NA | NA | NA | NA | 5 | 3358 | -6706 | 1316 | 0.000 | 0.64 | 0.05 |
| 0.057 | NA | -0.015 | NA | NA | NA | NA | NA | 4 | 3339 | -6671 | 1350 | 0.000 | 0.62 | 0.04 |
| 0.065 | NA | NA | -0.018 | NA | NA | NA | NA | 4 | 3314 | -6620 | 1401 | 0.000 | 0.66 | 0.01 |
| 0.047 | NA | NA | NA | NA | NA | NA | NA | 3 | 3293 | -6580 | 1441 | 0.000 | 0.65 | 0.00 |

**Table S10 Linear regression models of vulnerabilities of vertebrates from the Naracoorte ecological community. Vulnerabilities were calculated using the simulation method, and the table includes coefficient estimates, AICc and AICc weights, and adjusted $r^2$**

| intercept | basal connections | diet breadth | trophic level | basal connections* diet breadth | basal connections* trophic level | diet breadth* trophic level | basal connections* diet breadth* trophic level | df | logLik | AICc | delta | weight | adjusted $r^2$ |
|---|---|---|---|---|---|---|---|---|---|---|---|---|---|
| 0.281 | -0.173 | -0.327 | -0.259 | 0.199 | 0.158 | 0.308 | -0.186 | 9 | 2088 | -4157 | 0 | 1.00 | 0.99 |
| 0.273 | -0.078 | -0.277 | -0.259 | -0.013 | 0.078 | 0.270 | NA | 8 | 1968 | -3919 | 238 | 0.00 | 0.98 |
| 0.255 | -0.093 | -0.251 | -0.234 | NA | 0.084 | 0.236 | NA | 7 | 1920 | -3825 | 332 | 0.00 | 0.99 |
| 0.279 | 0.009 | -0.361 | -0.261 | -0.021 | NA | 0.349 | NA | 7 | 1724 | -3434 | 724 | 0.00 | 0.91 |
| 0.250 | -0.006 | -0.329 | -0.219 | NA | NA | 0.302 | NA | 6 | 1681 | -3350 | 807 | 0.00 | 0.90 |
| 0.258 | NA | -0.298 | -0.237 | NA | NA | 0.278 | NA | 5 | 1618 | -3226 | 931 | 0.00 | 0.90 |
| 0.178 | -0.157 | -0.013 | -0.158 | 0.019 | 0.134 | NA | NA | 7 | 1581 | -3148 | 1009 | 0.00 | 0.96 |
| 0.192 | -0.144 | 0.003 | -0.186 | NA | 0.136 | NA | NA | 6 | 1553 | -3095 | 1063 | 0.00 | 0.95 |
| 0.186 | -0.137 | NA | -0.176 | NA | 0.129 | NA | NA | 5 | 1550 | -3089 | 1068 | 0.00 | 0.80 |
| 0.132 | -0.020 | -0.028 | -0.099 | 0.022 | NA | NA | NA | 6 | 1407 | -2801 | 1356 | 0.00 | 0.73 |
| 0.147 | -0.002 | -0.009 | -0.130 | NA | NA | NA | NA | 5 | 1390 | -2769 | 1388 | 0.00 | 0.93 |
| 0.153 | NA | -0.008 | -0.138 | NA | NA | NA | NA | 4 | 1388 | -2767 | 1390 | 0.00 | 0.79 |
| 0.157 | NA | NA | -0.148 | NA | NA | NA | NA | 3 | 1374 | -2741 | 1416 | 0.00 | 0.56 |
| 0.159 | 0.001 | NA | -0.151 | NA | NA | NA | NA | 4 | 1374 | -2741 | 1416 | 0.00 | 0.79 |
| 0.070 | -0.051 | -0.065 | NA | 0.055 | NA | NA | NA | 5 | 1339 | -2667 | 1490 | 0.00 | 0.77 |
| 0.049 | -0.018 | -0.032 | NA | NA | NA | NA | NA | 4 | 1233 | -2457 | 1700 | 0.00 | 0.77 |
| 0.033 | NA | -0.028 | NA | NA | NA | NA | NA | 3 | 1119 | -2231 | 1926 | 0.00 | 0.24 |
| 0.021 | -0.015 | NA | NA | NA | NA | NA | NA | 3 | 1117 | -2228 | 1929 | 0.00 | 0.24 |
| 0.010 | NA | NA | NA | NA | NA | NA | NA | 2 | 1059 | -2114 | 2044 | 0.00 | 0.00 |
